# A neuron type-specific microexon in *Ank3*/ankyrin-G modulates calcium activity and neuronal excitability

**DOI:** 10.64898/2025.12.12.693948

**Authors:** Shah Alam, Georgia Dermentzaki, David Cabrera-Garcia, Miao Li, Ruizhi Wang, Melissa Campbell, Ilaria Balbo, Brittany L. Phillips, Min Li, Jessica Estrada, Marianna Zazhytska, Yow-Tyng Yeh, Lia Min, Elizabeth Rafikian, Elizabeth Valenzuela, Brian Joseph, Tulsi Patel, Dmytro Ustienenko, Helene Lovett, Huijuan Feng, Xiaojian Wang, Susan Brenner-Morton, Chyuan-Sheng Lin, Clarissa L. Waites, Hynek Wichterle, Lizhen Chen, Mu Yang, Edmund Au, Marko Jovanovic, Stavros Lomvardas, Paul M. Jenkins, Rui Yang, Sheng-Han Kuo, Yueqing Peng, Guang Yang, Neil L. Harrison, Chaolin Zhang

**Author notes:** To whom correspondence should be addressed (CZ), (NLH), (GY), (YP). Equal contribution (Shah Alam, Georgia Dermentzaki, David Cabrera-Garcia, Miao Li, Ruizhi Wang).

## Abstract

Recent studies have revealed many alternative exons differentially spliced across diverse neuron types in the mammalian brain, but their links to neuronal physiology remain unclear. Here we characterize a deeply conserved microexon E35a in *Ank3* encoding ankyrin-G (AnkG), a multifaceted adaptor protein best known as a master organizer of the axon initial segment (AIS) and as a leading genetic risk factor for bipolar disorder. E35a is predominantly skipped in cortical glutamatergic neurons but included in cortical GABAergic neurons and cerebellar neurons, which is dictated by multiple neuronal splicing factors. In E35a-deletion mice we generated, interneurons show increased excitability and somatic Ca^2+^ activity, without disruption in AIS. Biochemical analyses suggest that E35a inclusion facilitates AnkG interaction with a protein complex involving inositol trisphosphate receptors (InsP3Rs) important for intracellular Ca^2+^ signaling. Alternative splicing therefore allows AnkG to modulate neuron type-specific excitability in addition to its ubiquitous pan-neuronal role in organizing the AIS.

## INTRODUCTION

Proper function of the mammalian brain relies on a complex assembly of hundreds of neuronal cell types with distinct morphological and electrophysiological properties that are building blocks of intricate neural circuits. This remarkable cellular diversity is specified by precise neuron type-specific gene regulatory programs ^1, 2^. In addition to transcriptional regulation, alternative splicing of precursor messenger RNA (pre-mRNA) generates multiple mRNA and protein isoforms from a single gene by removing introns from pre-mRNAs and connecting exons in different combinations, thus dramatically amplifying the genetic information encoded in the genomic DNA ^3^. Regulation of alternative splicing is particularly extensive in the brain. In recent years, global transcriptome profiling using bulk or single-cell RNA-sequencing (RNA-seq) has enabled the discovery of distinct splicing patterns in the brain compared with non-neuronal organs ^4–6^, in neurons compared with non-neuronal cells in the brain ^7, 8^, in central nervous system (CNS) neurons compared with peripheral sensory neurons ^9^, as well as across different developmental stages in the brain ^9^. More recently, we and others have also identified hundreds of alternative exons differentially spliced in different neuronal types using RNA-seq data derived from purified neuronal populations or large-scale single-cell RNA-seq (scRNA-seq) ^10–12^.

These include microexons with lengths ≤ 30 nt, which are frequently neuron-specific and can be dysregulated in neurodevelopmental disorders such as autism ^13, 14^. The functional roles of a few such exons in the nervous system have been extensively studied. One example is Neurexin 1 (*Nrxn1*) exon 20 (also known as alternatively spliced site 4, or SS4), which is specifically included in GABAergic interneurons but skipped in excitatory neurons in a manner that is dependent on neuronal activity ^15^. This alternative exon modulates excitatory synapse formation during brain development by altering the postsynaptic binding partners of Nrxn1 ^15–17^. Multiple neuron type-specific exons and their regulatory mechanisms have also been studied in *C. elegans*. Two RNA-binding protein (RBP) splicing factors UNC-75 and EXC-7, whose mammalian homologs are CELF and ELAVL proteins, respectively, regulate a differential splicing program between GABAergic and cholinergic neurons ^18^. Among these, depletion of a neuron type-specific isoform of unc-64/Syntaxin, a critical component in the SNARE complex that mediates synaptic vesicle fusion, led to movement coordination deficits. Despite these salient examples, it remains unknown how most neuron type-specific alternative exons contribute to cellular and organismal phenotypes.

The family of ankyrins, composed of ankyrin-R (*ANK1*), ankyrin-B (*ANK2*), and ankyrin-G (*ANK3*), encoded by homologous genes, are adaptor proteins with shared domain architecture that link membrane proteins to the spectrin/actin cytoskeleton ^19, 20^. In this study, we focus on ankyrin-G (AnkG), encoded by *ANK3*, a leading risk gene for bipolar disorder (BD) in humans that has additional associations with schizophrenia (SCZ) and autism spectrum disorder (ASD) ^21–24^. AnkG is best known as the key organizer of the axon initial segment (AIS) and nodes of Ranvier, where it clusters ion channels, transporters, and cell adhesion molecules to physically separate axonal and somatodendritic compartments and enable the initiation and propagation of action potentials ^25–30^. Alternative splicing in *Ank3* is well known to contribute to AnkG isoform diversity ^31^ (Extended Data Fig. 1). The best known alternative exon is a giant exon (exon 37), which is specifically included in neurons to produce a 480 kDa isoform localized to the AIS and required for AIS establishment ^32^. In a previous study, we demonstrated that exon 27a (Extended Data Fig. 1), a developmentally regulated exon that is skipped in adult neurons, is also critical for AnkG to accumulate in the AIS by modulating AnkG-spectrin interactions. Forced inclusion of this exon results in ectopic expression of the embryonic isoform in maturing neurons, causing impaired AIS formation and dramatic reduction in neuronal excitability ^33^. This exon also contributes to AIS plasticity through neuronal activity-dependent splicing ^34^. Regulation of most of these alternative exons appears to be pan-neuronal, which is consistent with the notion that the AIS is a characteristic subcellular structure shared by most neuron types.

Here we investigate a highly conserved, neuron type-specific microexon, *Ank3* exon 35a or E35a, of 27 nucleotides (nt) long, which is highly included in GABAergic interneurons, but is predominantly skipped in glutamatergic excitatory neurons in the adult mouse cortex. We determine the major splicing factors that regulate cell type-specific splicing of this microexon. Furthermore, using an E35a-deletion mouse model, we demonstrate that AnkG has acquired neuron type-specific functions outside the AIS through alternative splicing, allowing it to fine-tune the electrophysiological properties of different neuron types.

## RESULTS

### A deeply conserved, highly regulated neuron type-specific microexon in *Ank3*

We recently performed systematic analysis of neuron type-specific alternative splicing in adult mouse neocortex using scRNA-seq data with deep, full-transcript read coverage ^12, 35, 36^. In this analysis, we found differential splicing among different neuron types defined at multiple hierarchical levels, including 469 cassette exons that showed differential splicing between glutamatergic and GABAergic neurons, the two major neuronal classes in the neocortex (Fig. 1a). Among them is a 27-nt microexon in *Ank3* (denoted E35a hereafter) located between constitutive exons E35 and E36 and upstream of the giant exon E37 (Extended Data Fig. 1). *Ank3* E35a is highly included in GABAergic interneurons, but is predominantly skipped in glutamatergic excitatory neurons (percent spliced-in, PSI or Ψ = 0.65 vs. 0.14) (Fig. 1b and Extended Data Fig. 2a,b). Splicing of the exon is neuron-specific, as it is skipped in non-neuronal brain cell types (Fig. 1b) and in non-brain tissues of both mice and humans (Extended Data Fig. 3a,b). Examination of additional neuron types in different brain regions using published RNA-seq data also revealed a striking degree of neuron type-specific splicing, with low exon inclusion in hippocampal glutamatergic neurons and medium spiny neurons of the dorsal striatum, similar to cortical glutamatergic neurons, but high inclusion in hippocampal GABAergic neurons, dopaminergic neurons, cerebellar Purkinje and granule cells, and olfactory sensory neurons, similar to cortical GABAergic neurons (Fig. 1c and Extended Data Fig. 2c; see further details in Extended Data Table 1 and Methods). During neurodevelopment, the exon inclusion is initially high in young glutamatergic neurons, as observed during neuronal differentiation *in vitro*, and is then reduced to low levels in mature neurons, a trajectory consistent with the overall decrease in exon inclusion observed in bulk cortex tissue in both the mouse and human brains; in contrast, E35a inclusion is maintained at a high level in GABAergic neurons (Extended Data Fig. 4a-c and data not shown).

**Figure 1:**
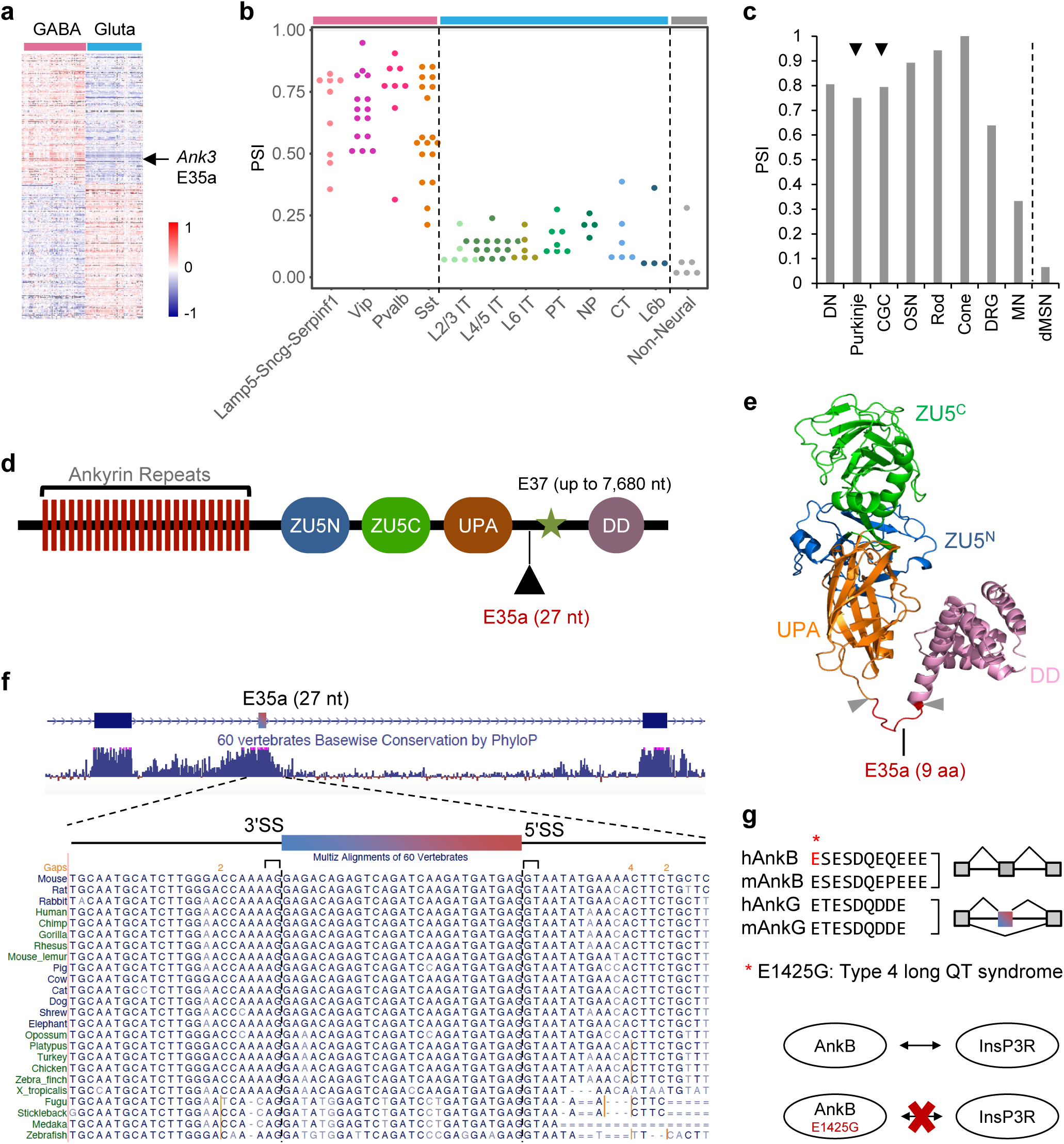
*Ank3* E35a shows GABAergic neuron-specific inclusion conserved in vertebrates. **a**, A heatmap showing the median-centered PSIs of cassette exons with differential splicing between GABAergic (magenta) and glutamatergic (blue) neuron types identified in adult neocortex scRNA-seq data ^35^. The different GABAergic and glutamatergic neuron subclasses were defined in the original study ^35^. *Ank3* microexon E35a (27 nt, between exon 35 and exon 36) is highlighted. **b**, E35a exon inclusion levels in individual transcriptional neuron types are shown. Cell types that belong to different neuronal classes, i.e., GABAergic neurons, glutamatergic neurons, and non-neuronal cells, are indicated using magenta, blue, and gray bars at the top, respectively. **c**, E35a exon inclusion levels in additional neuronal cell types in the adult mice. DN: dopaminergic neuron; CGC: cerebellar granule cells; OSN: olfactory sensory neuron; DRG: dorsal root ganglion; MN: motor neuron; dMSN: dorsal medium spiny neuron. Cerebellar neurons (Purkinje neuron and CGC) are highlighted by arrowheads since the relevant regions were also investigated in this study. **d**, A schematic showing AnkG protein domain architecture and the position of the peptide encoded by the microexon E35a. The position of the giant exon (E37, star) is also indicated. **e**, The structure of AnkG ZU5N/C-UPA-DD domain, as predicted by AlphaFold 3 (ref. ^38^). This structure is similar to the experimentally determined structure of its homolog, AnkB ^37^. Exon E35a, highlighted in red, encodes a 9-aa peptide in an intrinsically disordered region between UPA and DD domains. **f**, Conservation of E35a and flanking intronic regions across vertebrate species. 60-way conservation (phyloP) scores obtained from the UCSC genome browser are shown below the gene structure. Multiple alignments of E35a and flanking intronic sequences in a select subset of species are shown at the bottom. **g**, E35a is conserved between *Ank3* and *Ank2*, as shown in the alignment of amino acid sequences encoded by human and mouse *Ank3* E35a and its homologous exon in *Ank2*. Note that while *Ank3* E35a is alternatively spliced, the homologous exon in *Ank2* is constitutively included. An E1425G mutation in the peptide encoded by the *Ank2* exon is known to cause long QT syndrome. This mutation results in abrogation of protein-protein interaction between AnkB and inositol trisphosphate receptor (InP3R) and downstream alteration in intracellular calcium activity.

Multiple observations led us to focus on this exon as a model for detailed mechanistic and functional dissection of neuron type-specific alternative splicing. First, AnkG, as well as the other Ankyrin family members, is composed of 24 ankyrin repeats as a membrane-binding domain (MBD), two ZU5 domains and a UPA domain as a spectrin-binding domain (SBD), and a death domain (DD) followed by the C-terminal regulatory region. The peptide encoded by E35a is located in an intrinsically disordered region between the UPA domain and DD domain, based on the crystal structure of AnkB^37^ and the AlphaFold3-predicted structure of AnkG^38^ (Fig. 1d,e, and Extended Data Fig. 5a). This region is important for protein-protein interactions and is enriched in disease-related mutations^37^.

Interestingly, the inclusion of the nine amino acids encoded by E35a appears to convert a loop into part of an extended alpha helix upstream of the DD domain, based on AlphaFold3 prediction (Extended Data Fig. 5b). Second, E35a and its flanking intronic regions are highly conserved across vertebrate species, including humans and fish, which implies considerable functional significance and strong purifying selection to maintain the highly regulated splicing pattern over ∼450 million years of evolution (Fig. 1f). Third, to the best of our knowledge, the function of this microexon has not been characterized in the literature. However, when we searched for additional homologous sequences in the genome, we found a paralogous exon in the same region in *Ank2*, but not in *Ank1*. In contrast to the highly regulated alternative splicing for the *Ank3* microexon, the *Ank2* exon has six extra nucleotides, so that it encodes a total of 11 amino acids, and appears to be constitutively included in mouse tissues and cell types we examined (Fig. 1g). The homology between the *Ank2* and *Ank3* exons is also clear from amino acid sequence alignments (Fig. 1g). Interestingly, previous studies found that a mis-sense E1425G mutation in the homologous region in AnkB, which disrupts its interaction with inositol trisphosphate receptors (InsP3Rs), is associated with type 4 long QT syndrome and sudden cardiac death in humans ^39, 40^ (Fig. 1g). The deep conservation of AnkG E35a and the health relevance of the homologous region in AnkB suggest that it may play a role in delineating neuron type-specific physiological properties through highly regulated alternative splicing.

### Regulatory mechanisms of *Ank3* E35a alternative splicing

To study *Ank3* E35a, we first investigated which splicing factors contribute to the specific inclusion of the microexon in certain neuron types including GABAergic neurons. Our recent work identified multiple RBPs contributing to the differential splicing between glutamatergic and GABAergic neurons in the cortex, with Mbnl2, Celf2 and Khdrbs3 (Slm2) more highly expressed in glutamatergic neurons and facilitating glutamatergic neuron-specific splicing, while Elavl2 and Qk are more highly expressed in GABAergic neurons, facilitating GABAergic neuron-specific splicing ^12^ (Fig. 2a). By analyzing differential splicing upon genetic depletion of RBP splicing factors using RNA-seq datasets ^9, 41^, we identified at least four candidate RBPs regulating microexon E35a (Fig. 2b). In the embryonic brain, this exon is suppressed by Ptbp2. In contrast, Celf2 and nSR100 (Srrm4), which are more highly expressed in glutamatergic neurons and GABAergic neurons, respectively, activate inclusion of the exon. In the adult brain, a major splicing suppressor is Mbnl2, which is more highly expressed in glutamatergic neurons. E35a has a weak 3ʹ splice site (SS) immediately flanked by multiple purines (A/G) in the polypyrimidine tract (3ʹ SS: -0.98, 5ʹ SS: 7.96; based on the human homolog in VastDB ^42^). Accordingly, the upstream intron appears to be the main site of splicing regulation, which is also reflected in a high level of sequence conservation across vertebrate species. Indeed, the upstream intron contains binding sites of candidate RBP regulators we identified, based on CLIP data and bioinformatic motif site predictions ^9^ (Fig. 2c), suggesting that the regulation of E35a by these RBPs is direct and likely conserved during vertebrate evolution. Interestingly, we also found that splicing of E35a is neuronal activity-dependent through our reanalysis of published RNA-seq datasets ^43, 44^. Specifically, E35a inclusion is reduced in cultured cortical cells after depolarization induced by KCl treatment. In parallel, there is also an increase in Mbnl2 expression and a decrease in nSR100 activity (Extended Data Fig. 4d,e and ref. ^43^), which are consistent with, and presumably contribute to, the observed activity-dependent splicing change. Together, these data suggest that combinatorial regulation by multiple RBPs dictates the differential splicing of E35a between cortical glutamatergic and GABAergic neurons, which can be further modulated by neuronal activity.

**Figure 2:**
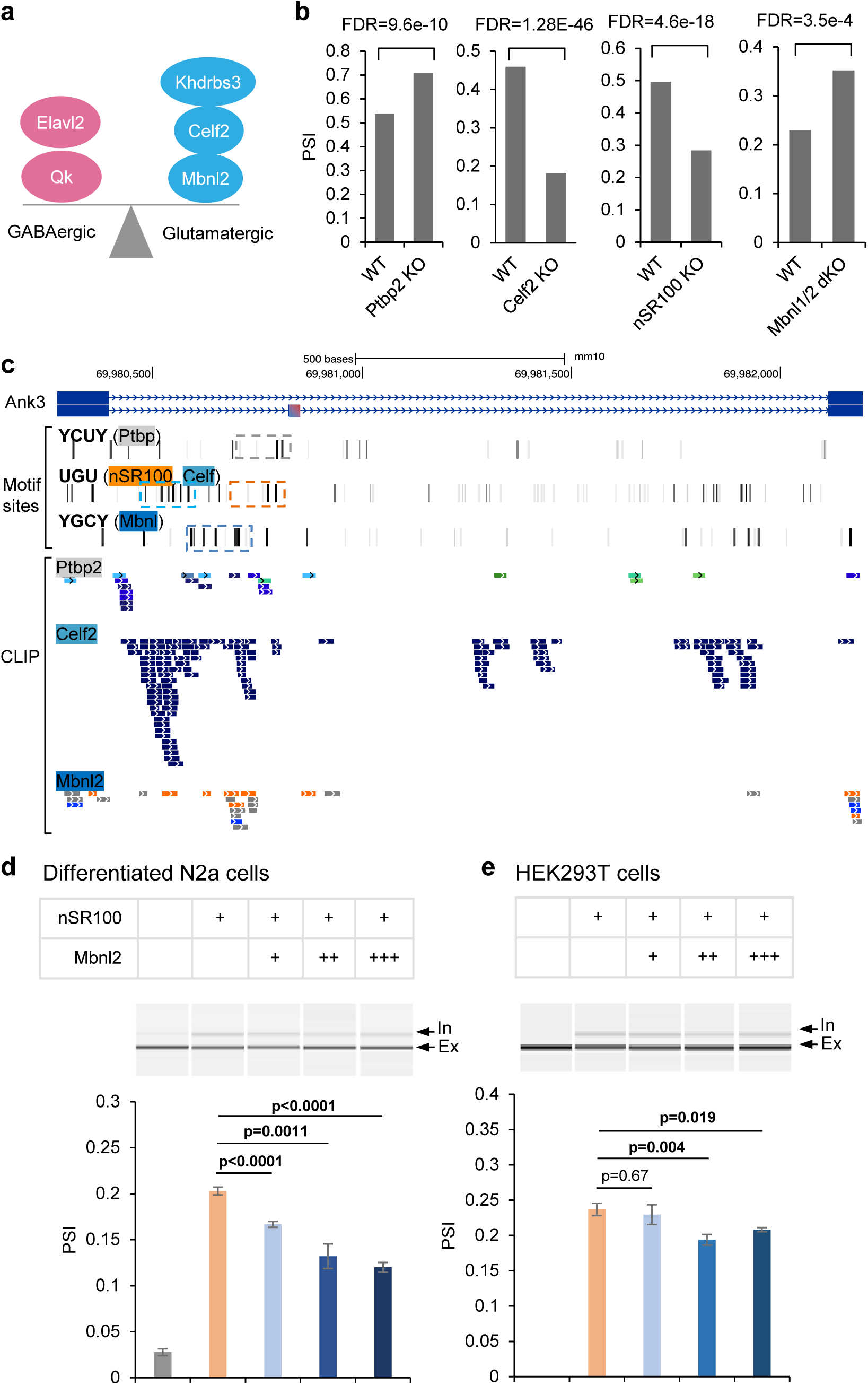
*Ank3* E35a splicing is under the combinatorial regulation of multiple neuron type-specific RNA-binding splicing factors. **a**, A schematic showing multiple RBPs that regulate GABAergic neuron-specific (Elavl2 and Qk) or glutamatergic neuron-specific (Khdrbs3, Celf2, and Mbnl2) splicing. **b**, *Ank3* E35a inclusion is suppressed by Ptbp2 and activated by Celf2, nSR100 (SRRM4), and Mbnl1/2, as determined by splicing changes quantified by RNA-seq in the mouse brain upon genetic depletion of each of the splicing factors. The statistical significance (false discovery rate or FDR) of differential splicing upon RBP depletion is indicated for each plot. **c**, Ptbp, Celf, nSR100, and Mbnl binding sites as evidenced by bioinformatically predicted motif sites and CLIP data. **d, e**, Validation of *Ank3* E35a splicing regulation by nSR100 and Mbnl2 using a minigene splicing reporter assay in differentiated N2a cells (d) and HEK293T cells (e). In each panel, a representative capillary electrophoresis (QIAxcel system) image is shown, with exon inclusion and exclusion bands indicated. nSR100 and Mbnl2 overexpression conditions are indicated at the top. The barplot at the bottom shows the quantification of exon inclusion (mean +/- SEM; n ≥ 5 per group). A two-sided t-test was used to evaluate the statistical significance for each pairwise comparison.

To further validate our results obtained from RNA-seq analysis, we focused on nSR100 and Mbnl2 as likely major regulators in mature neurons. We generated a mini-gene splicing reporter using sequences encompassing E35, E35a, E36, and intronic sequences in between. When the mini-gene was transiently transfected into differentiated mouse neuroblastoma N2a cells, E35a was completely skipped, similar to the splicing pattern of the endogenous *Ank3* in non-neuronal cells and tissues. Overexpression of nSR100 with the mini-gene led to an increase in exon inclusion to 20%, suggesting that nSR100 can effectively activate exon inclusion (Fig. 2d). Moreover, simultaneous overexpression of Mbnl2 together with nSR100 reduced exon inclusion to 12%, as compared to nSR100 expression alone, and the magnitude of Mbnl2-regulated splicing repression was dose-dependent (Fig. 2d). We also repeated this experiment in HEK293T cells and obtained similar results (Fig. 2e). The magnitude of Mbnl2-dependent skipping in HEK293T cells is less than that observed in N2a cells, probably because N2a cells more closely recapitulate the neuronal cellular context. Together, these cell-based experiments validated our results obtained from global transcriptome analysis using *in vivo* data derived from various mouse models.

### Generation of *Ank3* E35a deletion (E35a^-/-^) mice

To investigate the functional impact of E35a’s neuron type-specific splicing, we generated a mouse line lacking this exon using CRISPR-Cas9. Two single guide RNAs (sgRNAs) were designed to target the intronic sequences flanking E35a, resulting in precise deletion of the exon (Extended Data Fig. 6a,b).

Homozygous mutants, denoted *Ank3* E35a^-/-^, were successfully obtained using this approach (Extended Data Fig. 6c). RT-PCR analysis of RNA from the cortex and cerebellum confirmed that these mice expressed only the *Ank3* E35a-skipping isoform, while both isoforms were detected in wild-type (WT) controls; a higher exon inclusion level was observed in the cerebellum than the cortex, as expected from the major neuron types populating these brain regions (Extended Data Fig. 6d). *Ank3* E35a^-/-^ mice were born according to the expected Mendelian ratios and showed superficially normal development. Body weight assessment of adult *Ank3* E35a^-/-^ mice at 3 months of age did not exhibit any notable changes compared with WT controls (Extended Data Fig. 6e).

### *Ank3* E35a^-/-^ GABAergic neurons show increased neuronal excitability without AIS disruption

With the establishment of E35a^-/-^ mice, we first investigated the impact of E35a deletion on GABAergic neurons at the cellular level, with an initial hypothesis that the mutant mice might exhibit impaired AIS function and defects in neuronal excitability. For this analysis, given the associations of AnkG with epilepsy and bipolar disorders ^45^, we focused on the orbitofrontal cortex (OFC), a prefrontal region implicated in decision making and social behavior ^46, 47^.

To selectively label GABAergic interneurons with tdTomato, we crossed *Ank3* E35a^-/-^ mice with Dlx6a-Cre;Ai9 mice expressing tdTomato. Whole-cell patch clamp recordings were performed in tdTomato^+^ OFC interneurons using brain slices from *Ank3* E35a^-/-^ and WT control mice (Fig. 3a). Current-clamp measurements of passive and active membrane properties from resting membrane potential (RMP) in each neuron suggested that there was an increase in input resistance for *Ank3* E35a^- /-^ interneurons, while values of RMP and capacitance were similar to those of WT interneurons (Fig. 3b-d). Action potentials (APs) were elicited at RMP using depolarizing current commands, and the excitability of interneurons in both groups was compared using rheobase values and the number of APs evoked by each current step. We found that interneurons from *Ank3* E35a^-/-^ mice were more excitable than WT interneurons, as reflected in reduced rheobase (Mann-Whitney *U* test, p = 0.0391) and increased firing rate (two-way ANOVA, p = 0.0129, n = 46 neurons for E35a^-/-^ and 48 neurons for WT) (Fig. 3e).

**Figure 3.**
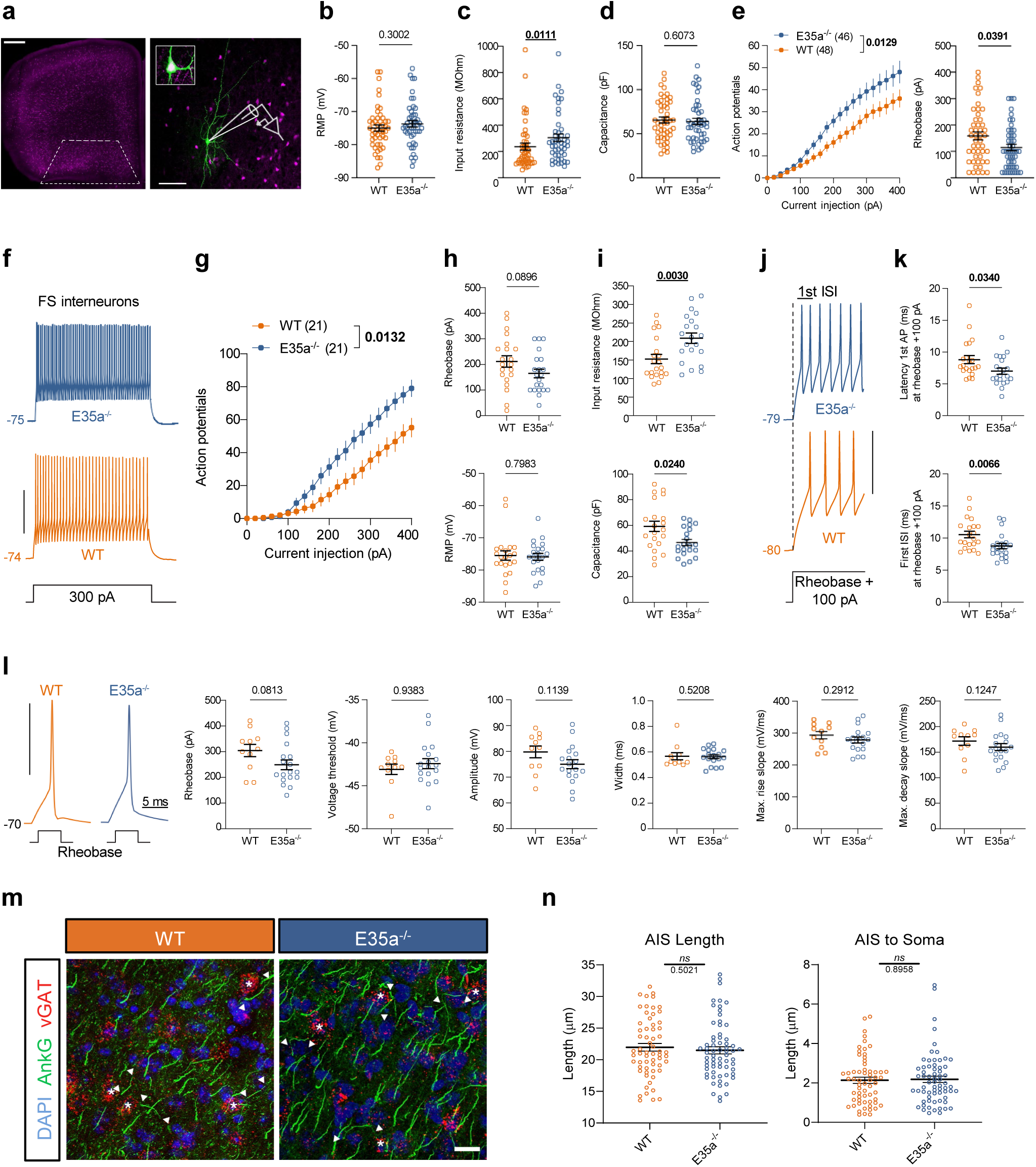
Intrinsic excitability is increased in fast-spiking *Ank3* E35a^-/-^ interneurons without apparent structural changes in the AIS. **a**, Illustrative confocal image showing *Dlx6a*-Cre:Ai9-tdTomato+ interneurons in a frontal coronal section with the OFC indicated by dashed lines (left panel). Scale bar: 500 µm. The right panel shows a schematic representation of a patch-clamp pipette and post-hoc immunostaining of a biocytin-filled interneuron during whole-cell recordings. Inset: enlarged section of the soma showing co-localization between filling (green) and Dlx6;tdTomato signals (magenta). Scale bar: 100 µm. **b-d,** Passive membrane properties in WT and *Ank3* E35a^-/-^ interneurons: resting membrane potential (RMP) (b), input resistance at RMP (c), and capacitance measured at RMP (d). **e,** Average number of action potentials (APs, left) and rheobase (right) during a series of 500-ms current injections from 0 to 400 pA in 20 pA incremental steps. Neurons were at resting membrane potential. Average data are presented as mean ± SEM. Note that the firing rate was higher (two-way mixed ANOVA, F_(1, 92)_ = 6.424, p = 0.0129) and rheobase lower (right panel, Mann-Whitney U test) in *Ank3* E35a^-/-^ interneurons compared to WT controls. **f,** Representative traces from the firing pattern of WT and *Ank3* E35a^-/-^ FS interneurons at RMP in response to a 500-ms stimulus with an amplitude of + 300 pA. RMP (mV) is indicated in each trace. Scale bar: 40 mV. **g,** Input-output curves show a higher number of evoked APs in Ank3 E35a^-/-^ interneurons compared to WT FS interneurons in response to increasing current step amplitudes (two-way ANOVA, F_(1, 40)_ = 6.725, p = 0.0132). Average data are presented as mean ± SEM. **h**, Rheobase (top panel) from data in panel (g), and RMP (bottom panel; WT: 75.5 ± 1.4 mV vs E35a^-/-^: 75.9 ± 1.1 mV). **i**, Input resistance at RMP (top panel; WT: 153 ± 12 MOhm vs E35a^-/-^: 209 ± 14 MOhm), and capacitance (bottom panel; WT: 59 ± 4 pF vs E35a^-/-^: 47 ± 2 pF) measured at RMP. **j**, First 35 ms of electrophysiological traces from WT and Ank3 E35a^-/-^ FS interneurons at RMP, evoked by a 500-ms stimulus at rheobase +100 pA. RMP (mV) is indicated in each trace. Scale bars: 40 mV (vertical), 10 ms (horizontal). **k**, Latency to the first AP (top panel; WT: 8.8 ± 0.6 ms vs E35a^-/-^: 7.0 ± 0.5 ms) and first ISI (bottom panel; WT: 10.5 ± 0.5 ms vs E35a^-/-^: 8.8 ± 0.4 ms) was measured at rheobase + 100 pA. **l**, Properties of the first AP at a common voltage (−70 mV) evoked from rheobase with 5 ms current injections in FS interneurons (bottom panels). The following properties of the first AP were analyzed: rheobase, voltage threshold, amplitude, width at half-height, maximum rise slope, and maximum decay slope. Data from 29 FS interneurons (WT = 11, *Ank3* E35a^-/-^ = 18). Scale bar: 40 mV (vertical). **b-l**, For statistical analyses in panels (b-e), data were derived from 48 WT interneurons (8 mice) and 46 *Ank3* E35a^-/-^ interneurons (8 mice). For statistical analysis in panels (f-k), data were derived from 21 WT interneurons (6 mice) and 21 *Ank3* E35a^-/-^ interneurons (7 mice). A mixed-model two-way ANOVA was used in panels (e, left panel) and (g). Mann Whitney U test was used in panels (b-d, e, right panel) and (h-l). Each dot represents data from a single cell. Data are presented as means ± SEM. Exact P-values are indicated in the graphs (p < 0.05 are in bold). **m**, Representative double-staining (FISH and IHC) image of AIS structures in vGAT+ GABAergic interneurons of WT and *Ank3* E35a^-/-^ mouse cortex. Arrowheads indicate AIS that can be clearly associated with vGAT+ cells selected for quantification. Scale bar: 20 µm. **n,** Quantification of AIS lengths (left) and AIS-to-Soma distances (right) in OFC vGAT+ GABAergic neurons. The numbers of AIS related measurements (61 AIS from 6 WT, and 66 AIS from 6 *Ank3* E35a^-/-^ mice) are indicated in the graph. No significant differences were identified in the t-test results of AIS length and AIS-to-Soma distance in GABAergic neurons between *Ank3* E35a^-/-^ and WT mice. All data are presented as mean ± SEM. By Mann-Whitney *U* test. ns indicates not significant (p>0.05).

Interneurons in the cortex consist of diverse neuronal subtypes with distinct firing properties ^2, 48–50^. To refine our analysis and reduce the variability in electrophysiological parameters due to interneuron diversity, we divided tdTomato^+^ interneurons into two groups: fast-spiking (FS; n = 42) interneurons and non-fast-spiking (non-FS; n = 52) interneurons. This categorization was based on the characteristic faster membrane time constant and higher firing rate of FS interneurons (Extended Data Fig. 7a) ^51–53^. We also confirmed that the age distribution of recorded neurons did not differ significantly between WT and *Ank3* E35a^-/-^ mice (Extended Data Fig. 7b), and that FS interneuron excitability was comparable between late adolescent and adult mice in both groups (Extended Data Fig. 7c).

We found that FS interneurons (n = 21 for E35a^-/-^ and 21 for WT) exhibited a higher firing frequency in *Ank3* E35a^-/-^ than WT (500 ms at 400 pA; 79 ± 5 vs. 55 ± 6 APs) (Fig. 3f, g), although the rheobase was not significantly lower in *Ank3* E35a^-/-^ neurons (Fig. 3h, top panel). RMP was similar between both genotypes (Fig. 3h, bottom panel), but FS interneurons from *Ank3* E35a^-/-^ mice displayed higher input resistance and lower capacitance, consistent with increased intrinsic excitability (Fig. 3i). APs evoked by rheobase +100 pA also showed shorter latency to the first AP and a decrease in the first inter-spike interval (ISI) in *Ank3* E35a^-/-^ (Fig. 3j,k), while the fast afterhyperpolarization (fAHP) was unaltered (Extended Data Fig. 7d). By contrast, non-FS interneurons showed no significant differences in firing properties between *Ank3* E35a^-/-^ and WT (Extended Data Fig. 7e, f), although the rheobase was lower in *Ank3* E35a^-/-^ cells (Extended Data Fig. 7g). The passive membrane properties of non-FS interneurons in *Ank3* E35a^-/-^ did not differ between genotypes (Extended Data Fig. 7h-j).

To further investigate the potential effect of the *AnkG* microexon on AP initiation related to the AIS, we analyzed the properties of the first AP evoked by rheobase stimulation ^50, 54, 55^. For FS interneurons, no differences were found in single AP properties elicited by either 5 ms current injections at a standardized membrane potential of -70 mV (Fig. 3l) or by 500 ms at RMP, except for the lower AP amplitude in *Ank3* E35a^-/-^ than in WT controls (Extended Data Fig. 7k). For non-FS interneurons, the properties of single APs were also similar between genotypes, whether evoked by long duration current steps at RMP (Extended Data Fig. 7l) or by short current pulses when neurons were held at -70 mV (Extended Data Fig. 7m). Taken together, these data show that *Ank3* E35a^-/-^ is associated with increased excitability in interneurons, specifically in presumed FS interneurons.

Given the altered neuronal excitability of FS cells, we examined AIS morphology to detect any potential structural changes ^56^. To our surprise, the comparison between *Ank3* E35a^-/-^ and WT mice did not show any apparent alterations in AIS length and distance of the AIS to the soma in cortical tdTomato^+^ interneurons within the OFC (Fig. 3m,n) or in other cortical regions such as primary motor and somatosensory areas that were also examined (data not shown). To further confirm this result, we transfected 480 kDa giant AnkG isoforms with or without E35a into dissociated hippocampal neurons from genetic *Ank3* knockout mice (*Ank3* ^-/-^) cultured *in vitro* (Extended Data Fig. 8a). Quantification of AIS length by immunostaining for AnkG or another AIS marker, β4-spectrin, at *in vitro* day 7 (DIV 7) did not reveal any notable differences (Extended Data Fig. 8b-d). Together, these results demonstrate that deletion of E35a increases interneuron excitability, particularly in FS subtypes, without detectable disruption of the AIS structure.

### *Ank3* E35a^-/-^ GABAergic neurons show increased somatic calcium activity

To investigate whether E35a contributes to neuron type-specific physiological properties *in vivo*, we performed two-photon Ca^2+^ imaging to examine the somatic activity of cortical neurons expressing the genetically encoded Ca^2+^ indicator GCaMP in awake, head-restrained mice ^57, 58^. This experiment was motivated by prior findings that a loss-of-function mutation in the homologous AnkB region causes aberrant Ca^2+^ handling in cardiac cells ^39, 40^. *Ank3* E35a*^-/-^* or WT control mice crossed with *Dlx6a*-Cre;Ai9 mice were injected with an adeno-associated virus (AAV) encoding GCaMP6s under the control of the synapsin promoter to drive indicator expression in both tdTomato^-^ pyramidal neurons and tdTomato^+^ interneurons (Fig. 4a). Three weeks post-injection, GCaMP6s-expressing neurons in the prelimbic (PL) area of the medial prefrontal cortex (mPFC) were imaged through a transcranial glass window (Fig. 4b). The PL region was chosen for accessibility for two-photon imaging; OFC and mPFC are reciprocally connected ^59^ and are both implicated in decision-making and social behavior ^46, 47^.

**Figure 4:**
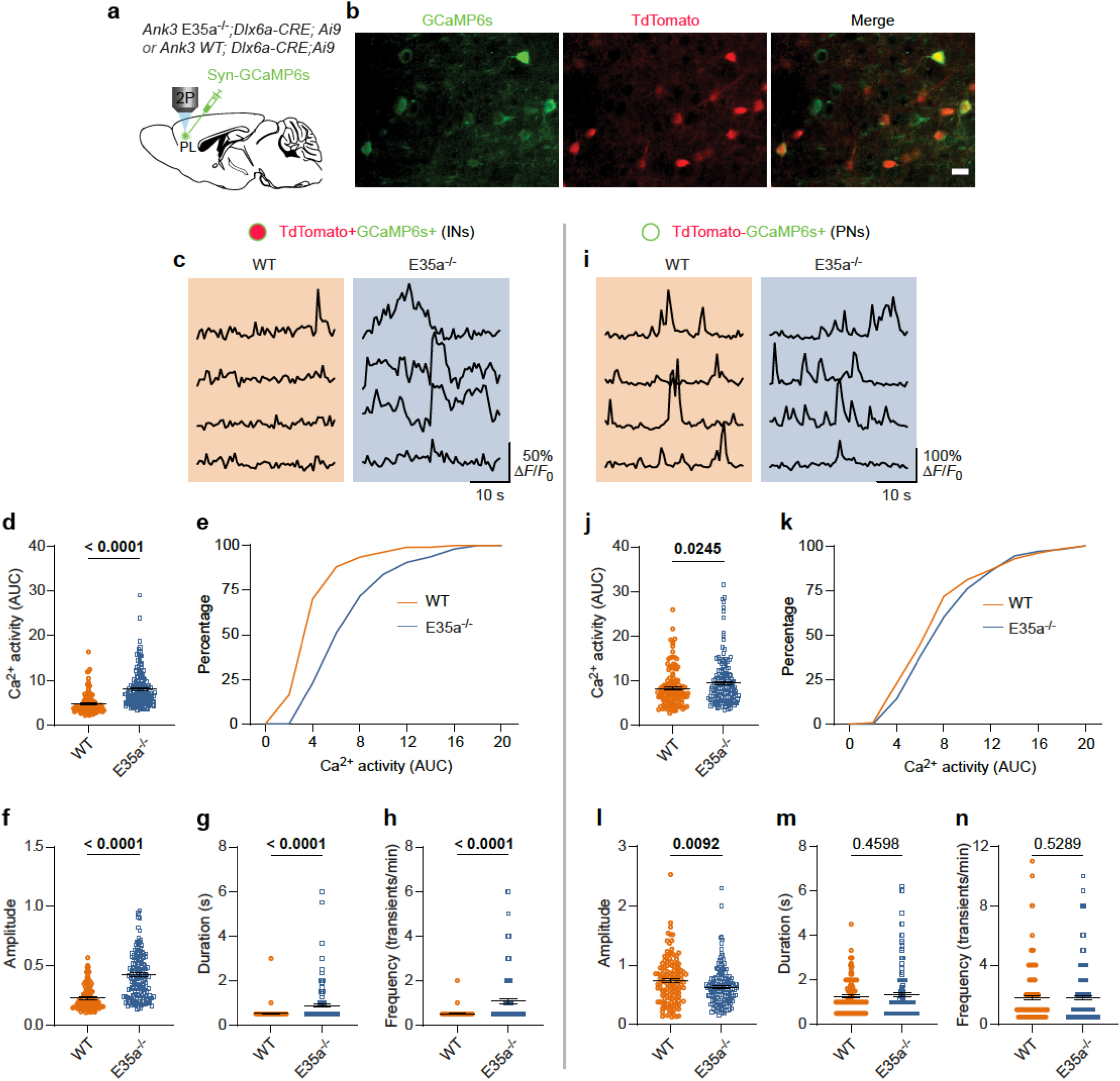
Increased interneuronal Ca^2+^ activity in *Ank3* E35a^-/-^ mice. **a**, Schematic of the experimental design: GCaMP6s was expressed in frontal cortical neurons via viral delivery, followed by *in vivo* two-photon (2P) imaging to examine neuronal activity. **b**, Representative images of GCaMP-expressing neurons in the cortex, with tdTomato-labeled interneurons (INs) highlighted. Scale bar: 20 µm. **c**, Representative Ca^2+^ fluorescence traces from neurons (Ins and PNs) co-labeled with GCaMP and tdTomato. **d**, Quantification of Ca^2+^ activity in interneurons, measured as the area under the curve (AUC) over 35 s (p < 0.0001). **e**, Cumulative distribution plot of AUC data shown in (d). **f**–**h**, Quantification of Ca^2+^ transient properties in interneurons: amplitude (f, p < 0.0001), duration (g, p < 0.0001), and frequency (h, p < 0.0001). **i**, Representative Ca^2+^ fluorescence traces from GCaMP-expressing pyramidal neurons (PNs), which lack tdTomato labeling. **j**, Quantification of Ca^2+^ activity in pyramidal neurons, measured as AUC over 35 s (p = 0.0245). **k**, Distribution plot of AUC data shown in (j). **l**–**n**, Quantification of Ca^2+^ transient properties in pyramidal neurons: amplitude (l, p = 0.0092), duration (m, p = 0.4598), and frequency (n, p = 0.5289). Throughout, each dot represents data from a single cell. In panels (d**–**h), *n* = 110, 164 cells from 3 mice for WT and 4 mice for *Ank3* E35a^-/-^ mice, respectively. In (j**–**n), *n* = 129, 170 cells from 3, 4 mice for Ank3 WT and *Ank3* E35a^-/-^, respectively. Data are presented as means ± SEM. Mann-Whitney *U* test was used to evaluate statistical significance (p < 0.05 are in bold).

We found that interneurons co-expressing GCaMP6s and tdTomato in *Ank3* E35a^-/-^ mice exhibited ∼2-fold higher somatic Ca^2+^ activity compared with WT controls during quiet resting states (Fig. 4c-h). Quantitative analysis revealed significant increases in area under the curve (AUC; 8.05 ± 0.31 vs. 4.75 ± 0.23, p < 0.0001), peak amplitude (0.42 ± 0.02 vs. 0.23 ± 0.01, p < 0.0001), duration (0.88 ± 0.06 vs. 0.53 ± 0.02, p < 0.0001), and frequency (1.08 ± 0.10 vs. 0.52 ± 0.01, p < 0.0001) of Ca^2−^ transients (*n* = 110 neurons from three mice for WT and 164 cells from four mice for E35a^-/-^; Mann-Whitney *U* test) (Fig. 4c-h). By contrast, pyramidal neurons (GCaMP6s^+^, tdTomato^–^) from *Ank3* E35a^-/-^ mice showed little change relative to WT (Fig. 4i). Minor differences were observed in AUC (Fig. 4j,k) and peak amplitude (Fig. 4l), whereas Ca^2+^ transient duration (Fig. 4m) and frequency (Fig. 4n) were unchanged (*n* = 129 WT neurons from three mice and 170 *Ank3* E35a*^-/-^*neurons from four mice; Mann-Whitney *U* test) (Fig. 4c-h). Together, these findings demonstrate that E35a deletion primarily increases somatic Ca^2+^ activity in cortical interneurons. This is consistent with the neuron type-specific splicing pattern of the microexon and with our *in vitro* electrophysiological results.

### E35a modulates protein-protein interactions with the NCX/NKA/InsP3R complex

It was previously shown in cardiac cells that AnkB can tether Na^+^/K^+^ ATPase (NKA) and Na^+^/Ca^2+^ exchanger 1 (NCX1) localized to T-tubules, invaginations of the plasma membrane (PM), and InsP3Rs in sarcoplasmic reticulum (SR) membrane to form a complex clustered into microdomains at SR-PM junctions. This complex plays an essential role in modulating cytoplasmic Ca^2+^ levels that are required for normal rhythmic contraction ^40^. Interestingly, the E1425G mutation in AnkB that causes type 4 long QT syndrome disrupts the NCX/NKA/InsP3R complex and the localization of its components at Z-line/T-tubules, resulting in altered Ca^2+^ signaling in cardiomyocytes ^39, 40^. We therefore tested whether alternative splicing of *Ank3* E35a similarly regulates Ca^2+^ activity in neurons by modulating protein-protein interactions.

We performed co-immunoprecipitation (co-IP) of type 1 InsP3R (InsP3R1) from mouse cerebellum extracts using an anti-AnkG antibody. The cerebellum was chosen over cortex because of the high E35a inclusion at the bulk tissue level in the former but not the latter (Extended Data Fig. 6d), reflecting major neuron types constituting these different brain regions (i.e., both granule cells and Purkinje cells in the cerebellum have high E35a inclusion, whereas the GABAergic neurons with high E35a inclusion represent a minority population in the cortex; Fig. 1b,c). Among the three InsP3R isoforms, InsP3R1 is the predominant brain subtype, forming micro-domains at the junctional region of the endoplasmic reticulum (ER; the equivalent in non-muscle cells of SR) and PM (ER-PM junctions; the equivalent of SR-PM junctions in non-muscle cells) ^60, 61^. Western blotting of the precipitated proteins using anti-AnkG antibodies identified three bands at approximately 190 kDa, 270 kDa, and, at a lower abundance, ∼480 kDa; the expression of the AnkG 190 kDa isoform was used for quantification in our analysis. An anti-InsP3R1 antibody detected a band corresponding to InsP3R1with a molecular weight of 268 kDa. In input samples, the abundance of InsP3R1 and AnkG are comparable between *Ank3* E35a^-/-^ and WT mice, suggesting that E35a deletion does not affect their expression levels. Importantly, in the co-IP samples, the amount of InsP3R1 pulled down from the *Ank3* E35a^-/-^ cerebellum showed a 77% reduction compared with that from WT (p = 0.003; n = 4 per group; paired t-test), confirming a reduced interaction between AnkG and InsP3R1 in the absence of the E35a microexon (Fig. 5a). We also probed NKA α1 and observed a ∼66% reduction in interaction with the complex in the absence of the E35a microexon (p = 0.048; n = 3 per group; paired t- test; Fig. 5b).

**Figure 5:**
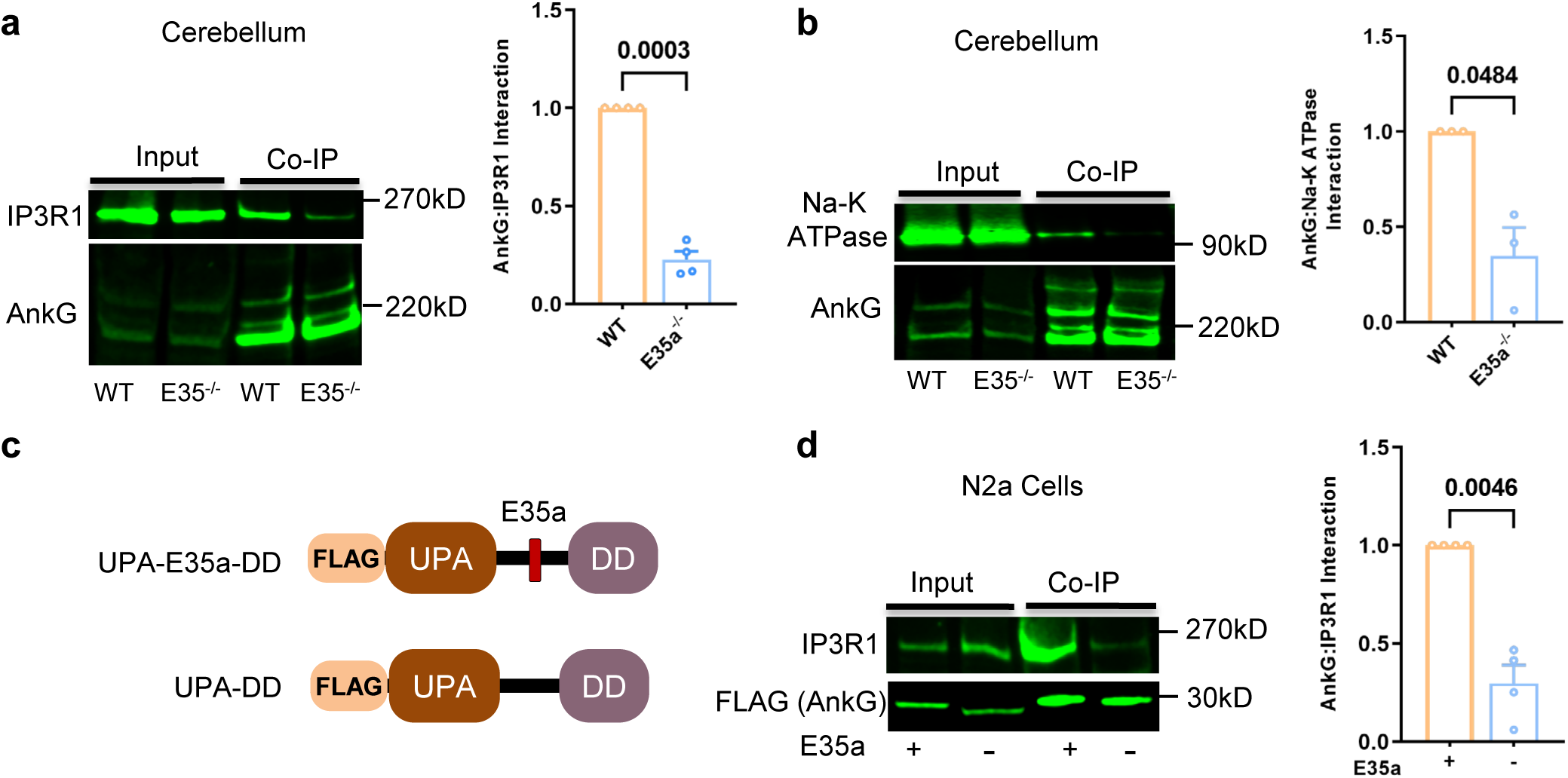
The absence of *Ank3* E35a abolishes AnkG binding to InsP3R1 and NKA. **a**, Relative binding of InsP3R1 (IP3R1) to AnkG in the WT and *Ank3* E35a^-/-^ mouse cerebellum. InsP3R1 from cerebellum lysate was co-immunoprecipitated using anti-AnkG and analyzed by immunobloting using anti-IP3R1. Representative immunoblot images are shown on the left and quantification is shown on the right. **b**, Similar to (a), but relative binding of Na-K ATPase to AnkG in the WT and *Ank3* E35a^-/-^ mouse cerebellum. **c**, Schematics of FLAG-tagged AnkG UPA-DD domain minigene with or without E35a. **d,** Relative binding of IP3R1, to the UPA-DD domains, with and without microexon E35a. UPA-DD minigene, with or without E35a, was transfected into N2a cells. IP3R1 from N2a total protein lysate was co-immunoprecipitated using anti-FLAG and analyzed by immunobloting using anti-IP3R1. For quantifications in panels (a, b, and d), data presented are mean +/- SEM. Relative binding in *Ank3* E35a^-/-^ or UPA-DD minigene without E35a was normalized to WT AnkG 190 kDa isoform (a, b) or UPA-DD minigene with E35a (d), respectively. n = 4 per group for panels (a) and (d) and n=3 per group for panel (b). Unpaired t-test with Welch’s correction was used to evaluate statistical significance (p < 0.05 are highlighted in bold).

To test whether the E35a-encoded peptide directly mediates these interactions, we generated FLAG-tagged mini-gene constructs encoding AnkG peptides encompassing UPA and DD domains, with or without the peptide encoded by the E35a microexon (Fig. 5c). Of note, since the full-length AnkG has a plethora of interacting proteins, the use of these constructs helps assess the specific impact of the microexon in mediating protein-protein interactions. The UPA-DD mini-gene, with or without E35a, was transfected into N2a cells, and co-IP of InsP3R1 was performed using the anti-FLAG antibody.

While the levels of the mini-gene and InsP3R1 expression were comparable in the input samples, the UPA-DD mini-gene without E35a pulled down only ∼30% of InsP3R1 relative to its counterpart containing E35a (p = 0.0046; n = 3 per group; paired t-test; Fig. 5d). This experiment corroborates our results *in vivo* indicating that the E35a-encoded peptide and its flanking protein domains can directly mediate protein-protein interactions between AnkG and the NCX1/NKA/InsP3R complex at ER-PM junctions involved in intracellular Ca^2+^ transport.

### Behavioral analyses in *Ank3* E35a^-/-^ mice

Given the altered AnkG isoform ratios, disrupted protein-protein interactions, and changes in neuronal excitability that we observed in Ank3 E35a^-/-^ mice, we next tested whether these molecular and cellular phenotypes manifest at the behavioral level (Extended Data Fig. 9a). Because E35a is highly included in cerebellar neurons, we first assessed motor function. Gait analysis revealed that *Ank3* E35a^-/-^ mice exhibited an expanded step length (p = 0.0004) and stance width (p = 0.0023), implying a broader base of support during locomotion (Extended Data Fig. 9b; n = 15 mice for WT and 14 mice for *Ank3* E35a^- /-^ group; Mann-Whitney *U* test; the same for other behavioral tests below). Front paw (FP) - hind paw (HP) distance was unchanged, but the alternation coefficient was decreased, indicating highly coordinated stepping with consistent left-right alternation in *Ank3* E35a^-/-^ mice. Mutant mice also crossed the testing platform faster than WT littermates (p = 0.0075) (Extended Data Fig. 9b). By contrast, beam walk performance was unaffected (Extended Data Fig. 9c). Collectively, these data suggest that *Ank3* E35a^-/-^ mice do not have gross coordination deficits but instead display a pattern of fast, frenetic locomotor activity.

Because *ANK3* variants are genetically linked to bipolar disorder ^21^, we also assessed the mice with assays of anxiety-related responses (open field and elevated plus maze; Extended Data Fig. 9d-e), memory (Y-maze, Extended Data Fig. 9f), and social behavior (3-chamber social, Extended Data Fig. 9g). No significant differences were observed between the WT and *Ank3* E35a^-/-^ mice in these assays. However, mutants showed a trend toward increased velocity in the 3-chamber test and more vertical counts in the open field, consistent with the hyperactive locomotor phenotype detected by gait analysis. Overall, the behavioral phenotypes of *Ank3* E35a^-/-^ mice are relatively mild.

## DISCUSSION

Technological advances in transcriptome profiling have led to remarkable progress in revealing the molecular diversity of different neuron types in the mammalian brain generated by alternative splicing, but the functional significance of this diversity is only just starting to emerge. Building on our previous systematic analysis ^12^, this study focuses on the regulation and function of a microexon in *Ank3*/AnkG, which is best known as a master regulator of the AIS. This exon is shared between *Ank3* and *Ank2* (but not present in *Ank1*), likely as a result of gene duplication, but the alternative splicing pattern appears unique to *Ank3* due to subsequent gene-specific adaptations after gene duplication followed by its fixation in vertebrates during over 400 million years of evolution. We found that combinatorial regulation by multiple splicing factors, including Ptbp, Celf2, nSR100, and Mbnl2, underlies its specific inclusion in GABAergic neurons and its skipping in glutamatergic neurons in the adult cortex. In the context of neurodevelopment, our analysis suggests a model in which E35a is initially suppressed by Ptbp in non-neuronal cells and very young neurons, and then activated by Celf2 and nSR100 in glutamatergic and GABAergic neurons, respectively. During neuronal maturation, the exon is repressed by Mbnl2, more so in glutamatergic than in GABAergic neurons, due to differential Mbnl2 expression, resulting in GABAergic neuron-specific exon inclusion in the adult cortex. This model is consistent with the major splicing factors that we identified by global analysis of neuron type-specific splicing ^12^. Moreover, the role of nSR100 in regulating neuron-specific splicing of microexons has been well documented ^13^. This model is also consistent with the decreased E35a inclusion upon neuronal activation, during which Mbnl2 expression increases and the activity of nSR100 decreases (Extended Data Fig. 4d,e and ref. ^43^). We tested and validated the role of Mbnl2 and nSR100 in E35a splicing using a splicing reporter assay in human and murine cells, although this assay did not quantitatively recapitulate *in vivo* splicing regulation in neurons; the clear but relatively modest effect of Mbnl2 is probably due to the involvement of additional co-factors. Indeed, the overexpression of Mbnl2 in mouse embryonic stem cell (ESC)-derived motor neurons induced a much stronger activation of E35a inclusion (Joseph et al., data not shown). However, it is important to note that while there is a good correlation between the splicing pattern of the exon and neurotransmitter types in the cortex and hippocampus, this is not always true in other brain regions. For example, cerebellar granule cells are glutamatergic but have a high level of E35a inclusion, whereas striatal medium spiny neurons are GABAergic ^62^ but have a low level of exon inclusion. Therefore, the neuron cell type-specific splicing of the exon is unlikely to be dictated simply by neurotransmitter phenotype, but rather by other mechanisms such as developmental lineages of neuron types.

Ankyrins are scaffolding adaptor proteins that link membrane proteins to the actin cytoskeleton in a variety of cell types ^20^. Given the instrumental role played by AnkG in neuronal physiology via the clustering of ion channels and cell adhesion molecules on neuronal and intracellular organelle membranes ^26, 27, 55^, we wondered whether the expression of different splice isoforms might enable it to acquire neuron type-specific functions. To this end, we generated a mouse model in which E35a is genetically deleted, so that these mice express only the E35a-skipping isoform. We have presented here a comprehensive survey of physiological, morphological and behavioral data relevant to neuronal excitability in mice harboring the *Ank3* E35a^-/-^ mutation.

Given the genetic link between AnkG and human bipolar disorder and the distinct splicing patterns of E35a, we decided to focus on prefrontal areas for most of our analysis, including electrophysiological and *in vivo* optical imaging studies. At the outset of this project, there was an expectation that this mutation might disrupt the structure and function of the AIS, and hence alter the generation and conduction of APs. However, the first conclusion that can be reached from our data is that this is clearly not the case. In GABAergic neurons of OFC within the cortex, where the exon35a is predominantly included, we recorded normal APs in mice lacking the exon. This is consistent with the lack of observed morphological defects in the AIS, suggesting the structure and function of the AIS are largely preserved in these mice. Importantly, we did see a significant increase in firing frequency and a ∼30% increase in input resistance, accompanied by a decrease in rheobase, a decrease in time to the initial AP and a decrease in inter-spike interval, specific to the FS cells, but not non-FS cells, in the OFC of *Ank3* E35a^-/-^mice, indicating an increase in the intrinsic excitability of FS cells. The resulting increase in firing frequency of FS cells is important because of the role of the PV+ interneurons in controlling the timing and synchrony of cortical oscillations (e.g., gamma oscillations) that are believed to be relevant to a variety of behaviors involving cortical circuitry ^53, 63^. It is also important to note that although the prefrontal cortex continues to develop through late adolescence ^64^, the intrinsic electrophysiological properties of cortical interneurons are already mature by the sixth postnatal week ^65, 66^. Indeed, our data confirmed that excitability in FS interneurons was comparable between late-adolescent and adult mice in both genotypes.

We also used *in vivo* calcium imaging to assess the impact of the *Ank3* 35a^-/-^ mutation on cortical circuitry. While brain slice recordings provide precise measurements of intrinsic excitability, they lack the spontaneous and modulatory activity present in intact networks due to deafferentation in the slice. In contrast, two-photon calcium imaging in awake mice allows monitoring of activity across multiple neurons within intact circuits, providing additional insights into the activity of these cells under the influence of local and extrinsic inputs. Using GCaMP6s to report on the fluctuation of free Ca^2+^ levels in identified neurons of the PFC, we showed that calcium activity in glutamatergic (pyramidal) neurons in *Ank3* E35a^-/-^ mice was largely normal. By contrast, GABAergic neurons showed a striking increase in activity, reflected in higher frequency and longer duration of Ca^2+^ transients. A role of AnkG in modulating Ca^2+^ activity is intriguing because calcium signaling governs many important cellular functions, including neurotransmitter release, synaptic plasticity, and neuronal excitability ^67–69^. Notably, another gene implicated in bipolar disorder is *CACNA1C*, encoding the alpha 1C subunit of the L-type voltage-gated calcium channel ^21^. Whether AnkG directly or indirectly interacts with CACNA1C is an interesting question that awaits further investigation.

At this point, we are uncertain about the origin of the changes in Ca^2+^ dynamics in FS cells of the *Ank3* E35a^-/-^ mice. Despite the complexity of the calcium dynamics in neurons, broadly speaking, Ca^2+^ may enter the cells via receptor-operated channels (ROCs) such as NMDA receptors or through voltage-dependent calcium channels (Ca_v_s). In addition, Ca^2+^ can be released from intracellular stores, which can control cellular Ca^2+^ dynamics ^67, 70^. An increase in the frequency of calcium transients in some GABAergic neurons of the PFC is certainly consistent with the increase in intrinsic excitability of FS cells recorded in brain slices, but could also reflect alterations in one or more other mechanisms. For example, intracellular Ca^2+^ can also activate SK and BK channels that contribute to the slow and fast AHP in several neuronal types ^71–73^. However, because no differences in fAHP were observed between genotypes, and fAHP in FS cells is primarily mediated by voltage-gated Kv3.1–Kv3.2 channels ^74–76^, the increased excitability of *Ank3* E35a^-/-^ FS cells is likely to be independent of Ca^2+-^gated BK/SK channels.

We investigated the possible biochemical basis underlying the observed changes in neuronal excitability and calcium activity in cortical GABAergic neurons of the E35a^-/-^ mice. A finding that initially helped motivate this study was that the AnkB peptide encoded by a paralogous exon of *Ank3* E35a is implicated in calcium signaling in cardiac cells. The AnkB peptide mediates the tethering of NCX and NKA to T-tubules and InsP3R on the sarcoplasmic reticulum (SR) membrane, resulting in the clustering of the NCX/NKA/InsP3R complex at SR-PM junctions. This complex facilitates the export of Ca^2+^ from the intracellular SR storage to the cell exterior, while a loss-of-function point mutation E1425G in AnkB results in the disassembly of the complex and elevated Ca^2+^ activity which causes a long QT syndrome in humans ^39, 40^. Analogous to the SR in cardiac myocytes, the endoplasmic reticulum (ER) in neurons forms a network of interconnected tubules and cisternae that extend from the outer nuclear membrane to the cytoplasm, axons and dendrites, and also contains numerous microdomains that are closely apposed to the plasma membrane (< 30 nm) or “ER-PM junctions” (see refs. in ^67^). Increasing evidence suggests that these ER-PM junctions play critical roles in regulating Ca^2+^ signaling in neurons. For example, a recent study demonstrated that ER-PM junctions are periodically distributed along the dendrites, interlinking a meshwork of ER tubules. These ER-PM junctions are populated with Ca_v_ channels and ER Ca^2+^-releasing ryanodine receptors (RyRs), and function as hubs for fine-tuning local Ca^2+^ homeostasis and facilitating the propagation of Ca^2+^ and electrical signals along the dendrites ^77^.

The homology between AnkG and AnkB led us to hypothesize that alternative splicing of *Ank3* E35a may allow neuron types with high inclusion of the exon, such as cortical GABAergic neurons, cerebellar Purkinje cells and granule cells, to tether the NCX/NKA/InsP3R complex and organize similar ER-PM microdomains to regulate calcium signaling as in cardiac muscle, but this capability might be absent or diminished in other neuronal types with low E35a inclusion. Consistent with this hypothesis, it was previously shown that the spectrin-binding domain of AnkG interacts directly with NKA ^78, 79^. Our co-IP experiments demonstrated that the inclusion of E35a is required for high-affinity interaction between AnkG and the NCX/NKA/InsP3R complex, as we demonstrated through comparison between WT and *Ank3* E35a^-/-^ cerebellar tissues or by the study of N2a cells overexpressing fragments of different *Ank3* isoforms (Fig. 5). We note that the direct assessment of AnkG-NCX/NKA/InsP3R interaction in the cortex is currently challenging, since GABAergic neurons with high E35a inclusion represent a minor population within the tissue and inclusion of the exon in the bulk cortex tissue is low. The disruption of the NCX/NKA/InsP3R complex in GABAergic neurons of *Ank3* E35a^-/-^ mice might result in reduced export of Ca^2+^ released from the ER storage, which would be consistent with the observed increased somatic Ca^2+^ activity. However, this model needs to be corroborated by direct observation of the complex at ER-PM junctions and its dependence on E35a inclusion. In addition, functional evidence for this proposed mechanism is currently lacking and awaits further investigation. Therefore, it remains unclear for now how, and to what extent, the ER storage mechanisms contribute to the observed Ca^2+^ activity changes, as compared to alternative mechanisms such as increased Ca^2+^ influx through ROCs or Ca_v_s upon neuronal depolarization.

Based on the pronounced changes in neuronal electrophysiological properties associated with E35a splicing, it was somewhat unexpected that the *Ank3* E35a^-/-^ mice did not show overt developmental abnormalities or major behavioral deficits. Across a battery of behavioral assays evaluating motor function, memory, anxiety- and depression-related behaviors, and social interaction, the only significant differences were increased step length and locomotion on gait analysis, consistent with a hyperactive phenotype. The divergence between the cellular and behavioral phenotypes may reflect compensatory adjustments in excitatory-inhibitory balance, particularly in behaviors requiring precise timing such as social interaction and memory. Alternatively, the mixed C57BL/6J and CBA/J (B6CBAF1) genetic background of the mice used here may have masked subtle behavioral deficits or introduced variability into the behavioral outcomes. Further studies using inbred C57BL/6J mice, with a focus on prefrontal cortex-dependent cognitive tasks, such as fear conditioning and extinction, will be important to fully assess the organismal impact of the inclusion or omission of this microexon.

In summary, our data provide compelling evidence for the functional significance of a neuron type-specific microexon in AnkG beyond its canonical role at the AIS, an aspect that was historically under-appreciated but has gained increasing interest and support in recent studies of neurons and other cell types ^80–84^. The neuron type-specific exon inclusion allows AnkG to acquire a highly regulated capacity to modulate calcium signaling proteins that demarcate neuronal membranes in certain cell types, and to fine-tune their electrophysiological properties through direct or indirect mechanisms. Our results also suggest the potential for parallel mechanisms between AnkB in cardiac cells and AnkG in neurons in membrane protein organization, which may represent another example of how the reuse and adaptation of existing molecular and cellular machinery underlie Darwinian evolutionary innovations that lead to the expanded phenotypic complexity of the nervous system in higher vertebrates, including mammals.

## ACKNOWLEDGEMENT

We thank members of the Zhang laboratory and George Mentis at Columbia University for scientific inputs and helpful discussions during this study, Columbia Transgenic Mouse Core Facility to produce *Ank3* E35a^-/-^ mice, and Columbia Neurobehavior Core Facility to assist behavior analyses. This study was supported by grants from the National Institutes of Health (NIH) (R01NS125018 to CZ, NLH and EA, R35 GM145279 to CZ; R01AA030604 to NLH; R35 GM131765 to GY, R01 NS136686, R01 NS118179, R01 NS124854 to SHK; R01NS080967 to CLW; R01AG070214 to LC; K99NS121136 to TP). Additional grant supports include Columbia RISE grant to CLW; Columbia Precision Medicine Initiative to YP; National Natural Science Foundation of China (32271016 to RY). BJ is a New York Stem Cell Foundation – Druckenmiller fellow.

## AUTHOR CONTRIBUTIONS

C.Z. conceived the study; S.A., G.D., M.C., and B.P. designed the experiments and led the project; M.Z., Y.-T.Y., B.J., T.P., H.F., and C.Z. contributed to analysis of splicing and gene expression using RNA-seq data; B.P. and C.-S.L. designed and generated *Ank3* E35a^-/-^ mice; S.A. and G.D. contributed to validation of *Ank3* E35a splicing regulation in brain tissues and cells using minigene reporters; G.D., M.C., I.B., J.E., E.R., and M.Y. contributed to behavior analyses; D.C.-G. contributed to patch-clamp recordings; R.W. and M.(Min) L., along with M.C., I.B., and B.P., contributed to immunostaining and AIS analyses; M.(Miao) L. contributed to calcium imaging; S.A., along with G.D., L.M. and E.V., contributed to analysis of AnkG interaction with its partner proteins. D.U., H.L., S.M. and L.C. contributed to reagent or resources; J.E. and X.W. provided technical assistance. C.Z. oversaw this study with C.W., H.W., M.Y., E.A., M.J., S.L., P.M.J., R.Y., Y.P., S.-H.K., G.Y., and N.L.H. supervising their respective lab members. G.D., S.A., D.C.-G., M.L., I.B., R.W., Y.P., R.Y., N.L.H., and C.Z. wrote the manuscript; All authors critically reviewed and approved the final manuscript.

## METHODS

### Mice

All experimental procedures followed the National Institutes of Health (NIH) Guide for Care and Use of Laboratory Animals (National Research Council, 2011). All animal procedures were approved and performed following the institutional animal care and use committee’s policies at Columbia University. Male and female mice were group-housed in polycarbonate cages with corncob bedding; they were maintained in a humidity-and temperature-controlled vivarium (20-22°C) on a 12/12 h light/dark schedule (lights on at 07:00 am and off at 07:00 pm). Animals had access ad libitum to food and water except during behavioral testing. *Ank3* E35a^-/-^ mice were generated in this study.

### Generation of *Ank3* microexon E35a deletion mice

To generate *Ank3* E35a deletion (E35a^-/-^) mice using CRISPR-Cas9, we designed two single guide RNAs (sgRNAs) flanking the microexon. The sgRNAs (IDT) were assembled with the Alt-R™ S.p. Cas9 Nuclease V3 (IDT) in an injection buffer containing 5mM Tris-HCl (pH 8) and 0.2mM EDTA. This assembly process aimed to form the CRISPR RNP (ribonucleoproteins) with a final concentration of 0.1 µM. The assembly was conducted at room temperature for 10 minutes. Subsequently, the assembled RNP was microinjected into the pronuclei of fertilized B6CBAF1 eggs. Eggs that survived the injection were then transferred into the oviducts of E0.5 pseudo-pregnant surrogates. Pups resulting from this injection were genotyped at the age of 15 days old (P15) using PCR. Heterozygous *Ank3* E35a^+/-^ mice were generated in a B6CBAF1 mixed background (Jackson Laboratory Stocks #100011) and bred to *Dlx6a*-Cre; Ai9^fl/fl^ mice (Jackson Laboratory Stocks #008199 and #007909).

### RNA-seq analysis

RNA-seq data used for *Ank3* E35a inclusion quantification in different neuron types were obtained from previous studies: cortical glutamatergic and GABAergic neurons, and non-neuronal cells from single-cell RNA-seq data ^35, 36^, hippocampal glutamatergic and GABAergic neurons ^10^, dopaminergic neurons ^85^, dorsal medium spiny neurons ^86^, cerebellar Purkinje cells and granule cells ^87^, olfactory sensory neurons ^88^, dorsal root ganglion (DRG) neurons ^89–91^, rod and cone cells ^92, 93^, and motor neuron ^94^. We also examined exon 35a splicing in mouse^95^ and human tissues^96^, during cortex development of mouse ^9^ and human brain ^97^, and differentiation of mouse embryonic stem cells (mESCs) to excitatory glutamatergic neurons ^98^. Most of these datasets were analyzed using Quantas ^7^ in a previous study ^9^ or otherwise analyzed using the same pipeline. Splicing across human tissues was calculated using exon junction read counts generated by the GTEx consortium ^96^. To identify potential regulators of exon 35a, we used RNA-seq data from mouse brains with genetic depletion of the following splicing factors, Ptbp2 (ref. ^99^), nSR100 (ref. ^41^), and Mbnl1/2 (ref.^9^). These datasets were also analyzed previously or in this study using the same pipeline ^9^. Conditional Celf2 knockout in cortical excitatory neuron were generated by breeding Celf2 ^fl/fl^ mice with *CamKII*-Cre, which was used to derive RNA-seq data of *Celf2* KO and WT control. The RNA-seq data was analyzed as described previously ^9^; only *Ank3* E35a inclusion was examined in this study, and additional analysis of the dataset will be described elsewhere. Neuron-activity dependent splicing was analyzed with the same pipeline ^9^ using RNA-seq data derived from primary cortical ^44^ and hippocampal ^43^ neurons depolarized by KCl. Public RNA-seq data analyzed in this study were summarized in Extended Data Table 1.

### Plasmid cloning

*Ank3* E35a minigene (genomic sequence extending from exon E35-E36 including E35a and flanking intronic sequences; 1,926 bp) was cloned from P7 mouse cortex genomic DNA in a TOPO TA vector (Thermo Scientific #450030) according to kit instructions (see primer sequences in Extended Data Table 2). Next, this sequence was subcloned to the pcDNA3.1 (+) vector, with EcoRI digest to both TOPO TA vector (+minigene) and pcDNA3.1 (+), followed by ligation and transformation.

The giant AnkG 480 kDa isoform construct without E35a (AnkG480-GFP) was described in our previous studies ^32, 100^. Due to the relatively large coding sequence (approximately 14K) of giant AnkG, we generated AnkG480-E35a-GFP including E35a using a cloning based strategy. In brief, the AnkG480-GFP plasmid was digested with different restriction enzymes, NheI and Bstz17I (Extended Data Table 2) to remove a fragment that contains the target region to insert E35a, and the cut vector fragment was purified. Fragment 1 was created by a forward primer that contains a 15bp overlap with the digested vector and a reverse primer that contains the mutated site and 15 bp overlap with fragment 2. Fragment 2 was created using a forward primer that contains the mutated site and 15 bp overlap with fragment 1 and a reverse primer that contains 15 bp overlap with the digested vector. Finally, the digested vector, fragment 1 and fragment 2 were assembled using In-Fusion® clone kit (Takara Bio USA, Inc.; #638920). Primers used to generate giant AnkG mutations are listed in Extended Data Table 2.

We generated plasmids expressing FLAG-tagged AnkG peptides containing specific UPA-DD domains with and without the E35a microexon under the control of the pCAGGs promoter. These sequences were cloned in the TOPO TA vector from cDNA derived from P7 mouse cortex with specific primers (Extended Data Table 2). These sequences were then subcloned into a pCAGGs plasmid together with an ATG start codon, 3X-FLAG, and stop codon sequences to allow for translation of these peptides.

### Cell cultures

HEK293T cells (ATCC #CRL-3216) were cultured in Dulbecco’s Modified Eagle’s Medium (DMEM; Gibco^TM^ #12491015) supplemented with 10% FBS (Thermo Fisher Scientific #A5256801), and 100U/ml penicillin-streptomycin (Thermo Fisher Scientific #15140122).

N2a cells (ATCC #CCL-131) were cultured in DMEM, supplemented with 10% bovine serum (Thermo Fisher Scientific #A5256801), 1x non-essential amino acids (Thermo Fisher Scientific #11140050), and 100U/ml penicillin-streptomycin. For N2a differentiation, cells were kept in the presence of 0.2% FBS and 2% DMSO (Sigma Aldrich # 34869) for 7 days.

### *Ank3* E35a minigene splicing reporter assay

HEK293T cells were plated at a density of 400,000 cells per cell in 6-well plates. Cells were co-transfected using calcium phosphate (CalPhos™ Mammalian Transfection Kit, #631312, Takara) with 125 ng *Ank3* minigene, 1 μg nSR100 (pLDPuro-MmnSR100N, Addgene #35171) and increasing amounts of MBNL2 (Addgene #217032; ref. ^9^) (1 μg, 1.5 μg, and 2 μg). N2a cells were plated at a density of 300,000 cells per cell in 6-well plates and differentiated for 7 days in the presence of 0.2% FBS and 2% DMSO. Cells were then co-transfected using lipofectamine 3000 (Thermo Fisher Scientific #L3000001) with 125 ng *Ank3* minigene, 0.5 μg nSR100, and increasing amounts of MBNL2 (0.5 μg, 1 μg, and 1.5 μg). Expression of transfected splicing factors was validated using western blots of RIPA cell lysates (data not shown) (Thermo Fisher Scientific #89901).

### RT-PCR validation of splicing quantification

To measure exon inclusion, RNA was extracted using TRIzol reagent (Invitrogen #15596026) or Direct-zol RNA kits (Zymo Research #R2061) from cultured cells or brain tissues. Total RNA was reverse transcribed using SuperScript III RT (Thermo Fisher Scientific #12574026,) in 20 μL reaction with random hexamer primers (f.c. 0.5 ng/μL). The cDNA was PCR amplified using gene-specific primers (Extended Data Table 2) and analyzed by the Qiaxcel system (Qiagen, #9001421) or agarose gel electrophoresis.

### AIS analysis using brain tissues

To analyze AIS of GABAergic neurons in *Ank3* E35a^-/-^ and WT adult mice (3-4 months old), we first labeled GABAergic neurons by *in situ* hybridization via the vesicular GABA transporter (vGAT) probe (SLC32A1, Advanced Cell Diagnostics #319191). Immunohistochemistry (IHC) was then performed to stain the AIS structure using anti-AnkG (Synaptic Systems #386003). Fluorescence *in situ* hybridization (FISH) was conducted using the RNAscope Multiplex Fluorescent Detection Kit v2 (Advanced Cell Diagnostics, #323110) following manufacturer protocol. 100μm coronal brain sections were cut using a vibratome (Leica VT1000s) from bregma 2.5 to 2.0 mm from 4% paraformaldehyde (Fisher Scientific #AAJ19943K2) perfusion-treated mice. The brain sections were then post-fixed with cold acetone for 2 minutes, followed by treatment with hydrogen peroxide, target retrieval reagent (Advanced Cell Diagnostics, #322000), and digestion with Protease III (Advanced Cell Diagnostics, #322337). The vGAT RNA probe (SLC32A1, Advanced Cell Diagnostics, #319191) was hybridized for 2 hours. Signal amplification was performed with HRP, followed by the fluorescent labeling with Opal 690 reagent (Akoya Biosciences, #FP1497001KT), which was diluted 1:1000 in TSA buffer (Advanced Cell Diagnostics, #322810). IHC was processed after the step of HRP blocker treatment in the FISH. The brain sections were blocked in blocking solution (5% donkey serum (Jackson ImmunoResearch, #017000121) with 0.3% Tween-20 in PBS solution) at room temperature for 1 hour, followed by incubation with anti-AnkG antibody (Synaptic Systems, #386003, 1:500 dilution in blocking solution) at 4°C overnight. Alexa Fluor 488 donkey anti-rabbit secondary antibody (Jackson ImmunoResearch, #711545152), diluted 1:500 in 3% BSA (Millipore Sigma, #12660) with 0.1% Tween-20 in PBS solution, was used to incubate the brain sections at room temperature for 1 hour.

Fluoro-Gel II (Electron Microscopy Sciences, #1798551), which contains DAPI, was used to mount the brain sections. Images were acquired using a Zeiss 810 confocal microscope with a 40x objective and Z-stack processing for the orbitofrontal cortical regions, the same brain region where our patch lamp analysis was performed. The positions of vGAT-labeled GABAergic neurons and their associated AIS structures were identified by observing different focal planes and 3D images from the Z-stack in ZEISS ZEN 3.8. The Extended Depth of Field (EDF) plugin (Biomedical Imaging Group) in ImageJ (FIJI) ^101^ was used to combine Z-stack images into a single, in-focus image ^102^. AIS length and AIS-to-Soma distance (from the root of the AIS to the soma boundaries) measurements were done by manual identification of the AIS region based on AnkG staining on EDF-treated images using ImageJ.

### AIS analysis using cultured hippocampal neurons

The method for culturing primary hippocampal neurons has been previously described in detail ^103, 104^. Hippocampi were harvested from genotyped postnatal day 0-1 (P0-P1) mice (AG^E22.23^ ^fl/fl^), then treated with trypsin, dissociated, and plated on poly-L-lysine-coated 18-mm glass coverslips at a density of approximately 600,000 neurons per 60-mm culture dish. The initial plating medium consisted of MEM supplemented with 0.6% (w/v) glucose and 10% (v/v) horse serum. After allowing the neurons to adhere for 2-3 hours, the coverslips were carefully transferred into preconditioned growth medium, which included Neurobasal-A medium with 2% (v/v) B27 supplement and 1×GlutMAX® (Thermo Fisher #21103049, #17504044, #35050061). Cultures were maintained in a 37°C incubator with 5% CO2.

#### Neuron transfection

In 3 DIV, 0.25 μg total AnkG plasmids (AnkG480-GFP and AnkG480-E35a-GFP) for each coverslip were mixed with 25 μL opti-MEM (Thermo Fisher, # 31985062) at room temperature for 5 minutes. Then the DNA mixture was combined with 0.75μL Lipofectamine 2000® (Thermo Fisher, #11668019) and 25μL Opti-MEM for 10 minutes at room temperature. Each coverslip was added 50μL of the DNA/lipofectamine mix and incubated for 30 minutes in the 37°C incubator. Then coverslips were flipped back to the previous neuron growth media. Transfected hippocampal neurons were maintained in the incubator until fixation.

#### Immunocytochemical staining

In the fixation step, we used incubate-warmed 4% wt/vol PFA with corresponding 4% wt/vol sucrose in 1x PBS to add on each coverslip and incubated for 10 min at 37 °C, and then washed with PBS three times for 5 minutes. For immunostaining, fixed neurons were permeabilized with 0.25% TritonX-100 for 10 min, washed with PBS three times for 5 minutes and then blocked with 5% BSA in PBS for 1 hour at room temperature. The primary antibodies (anti-AnkG or anti-beta4 spectrin, custom made rabbit polyclonal; ref. ^32^) were diluted (1:1000) in 5% BSA and incubated at 4°C overnight. On the next day, neurons were washed with PBS for 5 minutes and incubated with the appropriate fluorescent secondary antibodies in blocking buffer for 1 hour at room temperature. Finally, neurons were mounted with Mowiol mounting medium (Mowiol 4-88, Glycerol, Tris-HCl pH 8.5 and DABCO (1,4-diazabicyclo-[2,2,2]-octane) onto glass slides and allowed to cure for 24 hours before imaging.

#### Imaging and data analysis

Immunofluorescent stained neurons or paraffin-embedded tissue sections were imaged with confocal microscopy (Olympus FV3000). For AIS intensity quantification, images were captured using a 100x NA/1.4 oil objective. Each experiment was repeated independently three times, and the final quantification was derived from 8-10 neurons *in vitro*. To ensure the reproducibility of intensity measurements, all neurons from the same experiment were imaged under identical microscope settings. A standard AIS image was obtained as a three-step Z-series with a 0.30μm interval between each step. These Z-series images were processed using the maximum intensity projection function in Fiji software ^101^. The resulting maximum intensity projections were used for AIS intensity quantification, following the method adapted from Berger et al study ^105^. AIS length was quantified using a custom MATLAB script, similar as previously described ^100^.

### Electrophysiology

Brain slices were obtained from *Ank3* E35a^-/-^ (8) and WT control (8) mice (6 to 12 weeks old, except for one 15-week old mouse in each group) of both sexes. Mice were fully anesthetized with isoflurane and decapitated. Brains were dissected and sectioned in ice-cold (4˚C) artificial cerebrospinal fluid (aCSF) composed of (in mM): 124 NaCl, 2.5 KCl, 2 MgSO_4_, 1.25 NaH_2_PO_4_, 2 CaCl_2_, 26 NaHCO_3_, 10 glucose, and 2.5 sucrose, to yield coronal slices (300 µm thick) containing the orbitofrontal cortex (OFC) using a vibrating blade (VT1000S; Leica Biosystems). Slices were placed in a holding chamber containing oxygenated (95% O_2_-5% CO_2_) aCSF at 34 ˚C for 30 min and were subsequently transferred to room temperature for at least another 30 min before recordings were initiated. For recordings, slices were submerged in a chamber continuously perfused with oxygenated aCSF at 32 ˚C at a flow rate of 1-2 ml/min. Brain slices were visualized under an upright light microscope (BX51WI, Olympus) coupled to a camera (Hamamatsu #C8484), and the OFC area was identified according to cytoarchitectural criteria as previously defined in the Paxinos and Watson mouse brain atlas ^106^. *Dlx6a*-Cre interneurons were identified by the Ai9-tdTomato fluorescence (mercury arc lamp or LED 525 nm).

Whole-cell patch-clamp electrophysiology was performed as previously described ^107^. Intracellular recordings were made using pipettes pulled from borosilicate glass (World Precision Instruments, TW150F-4) with a resistance of 3-6 MΩ were filled with a potassium gluconate intracellular solution containing (in mM): 130 potassium gluconate, 4 NaCl, 10 HEPES, 0.3 EGTA, 4 Mg-ATP, 0.3 Na-GTP, 10 Na-phosphocreatine (300-305 mOsm). The pH was adjusted to 7.2-7.3 using KOH. 0.4% biocytin (Sigma #B4261) was added to the potassium gluconate intracellular solution for later morphological identification of a subset of recorded neurons. After recordings, slices were fixed for 24-48 h at 4 °C in 4% formaldehyde in PBS. After washing out the PFA, the slices containing biocytin-filled neurons were incubated with Streptavidin AlexaFluor-488 conjugate (1:300 v/v dilution, ThermoFisher # S11223) in PBS and Triton-X 0.3% for 3-5 h at room temperature. The slices were rinsed with PBS and mounted on glass slides to acquire images with a confocal microscope (Nikon Eclipse Ti) and then processed with Image J.

Recordings of the electrophysiological membrane properties and intrinsic excitability were performed in the absence of synaptic blockers. Electrophysiological recordings were performed with a MultiClamp 700B amplifier (Molecular Devices), and 1440 Digitizer (Molecular Devices), interfaced with Clampex 10.7 software (Molecular Devices). Data were collected at either 10 or 20 kHz and low-pass filtered at 2 or 10 kHz. Series resistance was compensated in current clamp using the bridge balance. Patch-clamp data was acquired only if the resting membrane potential was below -50 mV and initial access resistance <25 MΩ.

Following equilibration (∼2-3 min) after break-in, a current protocol of 10 pA command increments (6 steps starting at -30 pA, 500 ms) was used to measure passive membrane properties, including the resting membrane potential (RMP), the membrane time constant and capacitance. Input resistance (in MΩ) was calculated from the slope of a linear regression fit to the voltage-current relation in the 10 pA command increments. Using the same protocol, the time constant (tau) was estimated by the exponential fit between the beginning of the decay and the resting membrane potential. Then, capacitance was calculated from input resistance and time constant. Firing properties were characterized using 20 pA command increments from 0 to 400 pA, 500 ms duration, and APs were identified as events exceeding a threshold of -10 mV or with an amplitude of at least 70 mV from baseline. The input-output curves (the number of spikes in response to each current step) and rheobase (minimum current to elicit an AP) were determined using this protocol and the following parameters were also calculated for the first evoked AP: voltage threshold (voltage where dV/dt > 5% of the maximal increase dV/dt), amplitude (the difference between the AP voltage peak and the nadir of the fast afterhyperpolarization), duration (“width” at half-height), maximum rise slope (maximum dV/dt during depolarization phase), and maximum decay slope (maximum rate of fall of membrane potential during repolarization phase). Latency to the first AP (time between the start of the stimulus until the time of the first AP),first ISI (the ISI between the first two APs), and fAHP (voltage difference between voltage threshold and the minimum voltage after the AP peak), were measured at rheobase +100 pA. The 10 pA and 20 pA step protocols were repeated at RMP and from a holding potential of -70 mV. Additionally, single APs were evoked by short current pulses (5 ms) in 10 pA increments with the membrane potential held at -70 mV. AP properties using this protocol were analyzed as described above.

Offline analysis of *in vitro* electrophysiological data was performed using Clampfit 10.7 and Python scripts adapted from the IPFX package (https://github.com/alleninstitute/ipfx). To include neurons in the analysis, the following criteria were applied: cell with a stable resting membrane more negative than -55 mV, access resistance lower than 25 MΩ or variation <30% throughout the recording, input resistance lower than 1000 MOhm, and more than one AP evoked at RMP by the range of current stimuli (0 to 400 pA). Following these criteria, 5 WT and 3 *Ank3* E35a^-/-^ interneurons were excluded from the analysis. Graphing and statistical analyses were performed using GraphPad Prism 10 (GraphPad Software). Data are shown as box plots and as mean ± SEM in the input-output curves. Data was reported uncorrected for the liquid junction potential. Comparisons were considered significantly different when *p* < 0.05 and exact p-values were reported in the figures.

### *In vivo* calcium imaging

#### Surgical preparation

*In vivo* Ca^2+^ imaging was performed in the prelimbic (PL) area of the medial prefrontal cortex of awake, head-restrained mice, as previously described ^108, 109^. Mice were deeply anesthetized with an i.p. injection of 100 mg/kg ketamine and 15 mg/kg xylazine. After shaving the fur, the skull surface was exposed with a midline scalp incision, and the periosteum was removed. A head bar was glued to the skull with adhesive to minimize motion artifacts during imaging. A small skull region (∼1 mm in diameter) over the PL (anterior-posterior (AP) +2.68 mm, medial-lateral (ML) 0.5 mm) was carefully removed without damaging the dura mater. A round glass coverslip, matching the size of the removed bone, was glued in place, and dental acrylic cement was applied to secure the glass window. Throughout the procedure and recovery, the animal’s body temperature was maintained at ∼37 °C. After recovery, mice were habituated to head restraint on the imaging platform in three 10-min sessions to minimize stress. Imaging experiments were performed 24 h after surgery to ensure no lingering effects from anesthesia.

*In vivo Ca^2+^ imaging and data analysis*. GCaMP6s was expressed in PL neurons using recombinant adeno-associated virus (AAV) under the synapsin promoter (AAV1-Syn-GCaMP6s; Addgene #100843). 0.1 µl of diluted AAV (1:10) was injected into the left PL (AP +2.68 mm, ML 0.5 mm, subpial (SP) 0.85 mm) of *Dlx6a*-Cre;Ai9+; *Ank3* E35a^-/-^ or *Dlx6a*-Cre;Ai9+; *Ank3* WT mice. The injection was performed over 10–15 min using a Picospritzer III (15 p.s.i., 10 ms, 0.5 Hz) and a glass microelectrode. Imaging was performed approximately 3 weeks post-injection. *In vivo* Ca^2+^ imaging was conducted using a Scientifica two-photon system equipped with a Ti:Sapphire laser (Chameleon Vision-S, Coherent) tuned to 920 nm. Imaging was performed using a 25× objective (1.05 N.A.) immersed in artificial cerebrospinal fluid, with a digital zoom of 1.5×. Images were acquired at ∼1.7 Hz (2-μs pixel dwell time) with a resolution of 512 × 512 pixels. Data acquisition was performed using ScanImage software, and subsequent analyses were conducted using NIH ImageJ software. Regions of interest (ROIs) corresponding to visually identifiable somas labeled with GCaMP6, with or without tdTomato, were selected for quantification. The fluorescence time course for each ROI was determined by averaging pixel intensities within the ROI. Background fluorescence was subtracted from the GCaMP6s signal, and Ca^2+^ transients were calculated as Δ*F*/*F*_0_, where Δ*F*/*F*_0_ = (*F*–*F*_0_)/*F*_0_, with *F*_0_ representing the baseline fluorescence averaged over a 3.5-s period of minimal fluorescence during the recording. Motion artifacts due to respiration and heartbeat were generally <2 µm and were minimized by habituation and head bar fixation. Vertical movements were rare, and segments with struggling behavior were excluded from analyses.

#### Quantification and statistical analysis

The average integrated Ca^2+^ activity was quantified as the area under the curve (AUC) of Δ*F*/*F*_0_ over 35 s. Ca^2+^ events were defined as fluorescence signals > 3× the standard deviation (s.d.) of baseline fluorescence. The peak amplitude of Ca^2+^ transients was defined as the average of peak Δ*F*/*F*_0_ during the 35 s recording. Frequency was calculated as the number of Ca^2+^ transients per min, and duration was measured as the average full width of Ca^2+^ transients during the recording period.

Summary data are presented as means ± SEM. Sample sizes were chosen to ensure statistical power while minimizing animal usage. No successfully imaged animals were excluded from analysis. Data normality was assessed using the Shapiro-Wilk test. Nonparametric statistical comparisons were made using the Mann-Whitney *U* test, with significance set at p < 0.05. All statistical analyses were performed using GraphPad Prism software.

### Co-Immunoprecipitation

#### Brain tissue (Cerebellum)

Cerebellum lysates from WT and *Ank3* E35a^-/-^ mice brains were prepared by homogenizing the tissue using a hand Dounce in lysis buffer (30mM Tris, 150mM NaCl, 1% IGEPAL, 0.5% Na Deoxycholate, 0.1% SDS, 2mM EDTA, pH 6.8) with freshly added protease and phosphatase inhibitors. The homogenates were ultracentrifuged at 100,000 rpm for 30 minutes at 4°C to collect clear whole tissue lysates.

For the co-immunoprecipitation (co-IP) process, 50 μL of magnetic beads (Protein G, Invitrogen, #10003D) were prepared by resuspending them in a tube, separating them from the solution using a magnet, and washing them with NT2 buffer (50 mM Tris-HCl, pH 7.4, 150 mM NaCl, 1 mM MgCl₂, 0.05% NP-40). Subsequently, 5 µg of the anti-AnkG antibody (Santa Cruz #sc-12719) was added to the magnetic beads and incubated for 4 hours at room temperature. The supernatant was removed using a magnet, and the magnetic bead-antibody complex was resuspended in whole tissue lysates (500µg total protein) and rotated overnight at 15 rpm at 4°C. The following day, the bead-antibody-antigen complex was washed three times with NT2 buffer and once with wash buffer (NT2 buffer without NP-40), then resuspended in a clean tube. The target antigen was eluted from the complex using 50 µL elution buffer (50 mM glycine, pH 2.8) by resuspending the magnetic bead-antibody-antigen complex and placing it on a thermomixer at 23°C and 500 rpm for 15-20 minutes. The tubes were then placed on the magnet, and the supernatant was transferred to new tubes. To adjust the pH, 2 µL of Tris-HCl pH 7.5 was added along with 12.5 µL of premixed NuPAGE™ LDS Sample Buffer (Thermo Fisher Scientific #NP0007), and the samples were stored at -20°C. When ready for Western blotting, NuPAGE™ Sample Reducing Agent (Thermo Fisher Scientific # NP0009) was added to the samples, which were heated for 10 minutes at 68°C.

#### N2a Cells

N2a cells were cultured in DMEM GlutaMAX (Invitrogen #10566-016) supplemented with 10% fetal bovine serum. For the co-IP experiments, cells were plated on a 6-well plate, with three wells designated for each condition: with and without E35a. After 48 hours of transfection with minigene plasmids, cells from three wells of each condition were pooled together. The cells were lysed in 400 µL of IP lysis buffer (Pierce™ IP Lysis Buffer, Thermo Fisher Scientific #87787) with protease and phosphatase inhibitors and ultracentrifuged at 100,000 rpm for 30 minutes at 4°C. Total protein isolated from all three wells for each condition was used for IP using Anti-Flag M2 antibody (Sigma-Aldrich F3165), following the same protocol as described above for cerebellum samples.

#### Western immunoblotting

Protein lysates were separated by SDS-PAGE and transferred onto a nitrocellulose membrane using the iBlot Dry Blotting System (Thermo Fisher Scientific #IB21001). The membranes were blocked with 5% bovine serum albumin in TBS-T (50 mM Tris-HCl, pH 7.4; 150 mM NaCl; 0.1% Tween-20) for 30 minutes at room temperature. They were then incubated overnight at 4°C with the primary antibodies IP3R1 (Millipore Sigma #ABS55) and/or Na-K ATPase (Thermo Fisher Scientific # MA5-36259) (1:1000). Following incubation, the membranes were washed three times with TBS-T buffer (10 minutes each wash) and then incubated for 1 hour at room temperature on a rotator with an HRP-conjugated secondary antibody. The membranes were washed again three times (10 minutes each wash) to remove excess secondary antibody. Next, the membranes were incubated with infrared Dye 800-conjugated secondary antibodies (1:10,000 dilution, LI-COR Biosciences, #926-32210 and #926-32213) for 1 hour at room temperature. Images were captured using the Odyssey infrared imaging system (LI-COR Biosciences). Quantification was performed using ImageJ and Prism GraphPad software 9.0.

### Behavioral assays

All behavior analyses were performed using ∼3-month-old mice during the light phase of the light/dark cycle. All behavior tests, except for the 3-chamber test, were performed on 15 WT (9 males and 6 females) and 14 *Ank3* E35a^-/-^ (6 males and 8 females). Three-chamber test included 21 WT (11 males and 10 females) and 19 *Ank3* E35a^-/-^ (9 males and 10 females).

#### Gait analysis

Gait analysis was performed as previously described^110^. In brief, a transparent Plexiglass walkway (66 cm long, 6 cm wide) connected to a black box at its end was used to entice the animals in. The walkway has a mirror underneath with an inclination of 45° to allow a clear vision of the paws disposition from the front. Mice were filmed with a digital camera 12MP and 60 fps placed horizontally, 35 cm from the walkway. *Ank3* E35a^-/-^ and WT littermates were encouraged to walk at least three consecutive steps per trial, for a total of three trials. Still-frames from the recordings were extracted and analyzed offline using the Manual Tracking plugin of Fiji software ^101^ in order to obtain data concerning stride length (the distance of the same paw in a consecutive step), width (the distance between the center of the two hind or fore paws), the distance between ipsilateral fore paw and hind paw placements and the fore and hind stance (alternation coefficient ). The walking speed was also measured in order to exclude differences, which might affect gait parameters.

#### Beam walk test

The beam walk test was performed using a beam (55 cm long section, 5 mm wide) connected to a black box at its end to entice the animals in. Mice were trained in the beam one day prior to testing by allowing them to cross the beam for three consecutive runs. The test result was recorded using Noldus’ Ethovision software, which uses mouse body point tracking features while crossing the beam. Parameters such as Mean Velocity (cm/s), Mean Meander (deg/cm) and % Mean Mobility were analyzed. A camera was also used in parallel to assess manually the number or toe slips of each mouse.

#### Y-maze (spontaneous activity)

Spontaneous alteration is assessed in a Y-shaped maze with three white, opaque plastic arms at a 120° angle from each other. After introduction to the center of the maze, the animal is allowed to freely explore the three arms for a total of eight minutes. Over the course of multiple arm entries, the subject should show a tendency to enter a less recently visited arm. The number of arm entries and the number of triads are recorded in order to calculate the percentage of alternation. An entry occurs when all four limbs are within the arm. This test is used to quantify cognitive deficits in transgenic strains of mice. Percent alternation can be analyzed as a measure of working memory. Alternation%= (Number of spontaneous alternations)/(total number of arm entries-2) ×100, where one alternation refers to a sequence of three different arms. The theoretical chance alternation% = 1 × 2/3 × 1/3 ≍ 22%.

#### Three-chamber test

Social approach was assayed using an automated three-chambered apparatus (NIMH Research Services Branch, Bethesda, MD) as previously described ^111, 112^. Novel target mice were 3-months old FVB/NJ mice of the same sex as the subjects. The apparatus was a rectangular, three-chambered box made of clear polycarbonate. Retractable doorways built into the two dividing walls controlled access to the side chambers. Number of entries and time spent in each chamber were automatically detected by photocells embedded in the doorways and tallied by the software. The test session began with a 10-min habituation session in the center chamber only, followed by a 10-min habituation to all three empty chambers. Lack of innate side preference was confirmed during the second 10-min habituation. The subject was then briefly confined to the center chamber while the clean novel object (an inverted stainless-steel wire pencil cup; Galaxy; Kitchen-Plus; http://www.kitchen-plus.com) was placed in one of the side chambers.

A novel mouse previously habituated to the enclosure was placed in an identical wire cup located in the other side chamber (a disposable plastic drinking cup containing a lead weight was placed on the top of each inverted wire pencil cup to prevent the subject from climbing on top). After both stimuli were positioned, the two side doors were simultaneously lifted and the subject was allowed access to all three chambers for 10 min. Time spent in each chamber and entries into each chamber were automatically tallied. Time (frequency and duration) spent sniffing and rearing towards the novel object and time spent sniffing and rearing towards the novel mouse during the 10-min test session were later scored from video recording, by an observer using two stopwatches. The apparatus was cleaned with 70% ethanol and water between subjects. FVB/NJ mice were used as target novel mice because this strain is generally inactive, passive, and does not exhibit aggressive behaviors toward subject mice. Using a minimally active partner is a strategy that allows all approaches to be initiated by the subject mouse only.

#### Open field

Open field test was performed according to published protocols ^113, 114^. Each mouse was gently placed in the center of a clear Plexiglas arena (27.31× 27.31× 20.32-cm, Med Associates ENV-510) lit with dim light (∼5 lux), and was allowed to ambulate freely. Infrared (IR) beams embedded along the x, y, z axes of the arena automatically track distance moved, horizontal movement, vertical movement, stereotypies, and time spent in the center zone. Exploration was monitored during a 30-min session with Activity Monitor Version 7 tracking software (Med Associates Inc.). Parameters such as ambulatory distance (cm), center time (sec), and vertical counts were recorded. Data were analyzed in six, 10-min time bins. Areas were cleaned with 70% ethanol and thoroughly dried between trials.

#### Elevated plus maze (EPM)

The elevated plus maze (EPM) apparatus consists of two open arms (30 cm x 5 cm) and two closed arms (30 cm x 5 cm) extending from a central junction (5 cm x 5 cm) area. Each mouse completes one trial with a duration of 5 minutes. Each mouse is gently placed in the junction area to start the trial and is allowed to ambulate freely in the maze for the duration of the trial. Exploration of the closed and open arms as well as total entries were monitored via the Ethovision XT software. Percent (%) time spent in closed/open arm was calculated by dividing the time spent in closed/open arm by 300 seconds, and multiply by 100). Total entries are the sum of the open entrance and closed entrance. Areas were cleaned with 70% ethanol and thoroughly dried between trials.

### Statistical analysis

Statistical analysis was performed using GraphPad Prism unless indicated otherwise. Specific tests used were also provided as described in results or figure legends (see summary in Extended Data Table 3).

## Data availability

All data generated or analyzed during this study are included in this published article (and its supplementary information and Source Data files). RNA-seq data used in this study were obtained from published studies, as described in Methods (see summary in Extended Data Table 1).

## SUPPLEMENTARY FIGURE LEGENDS

**Extended Data Figure 1: ANK3 gene structure and alternative splicing isoforms.**

The human *ANK3* gene, as represented by the UCSC gene models, is divided into multiple segments, with overlaps shown between consecutive segments, to enhance visibility. Constitutive exons are indicated in black and alternative exons are labeled in blue. Not all alternatively spliced exons are shown. Protein domains encoded by specific exons are also indicated on the top of each segment.

**Extended Data Figure 2: Differential splicing of *Ank3* microexon E35a in diverse neuronal cell types.**

**a,** Quantification of E35a inclusion in individual glutamatergic and GABAergic neuron types in Tasic 2016 dataset. E35a exon inclusion levels in individual neuronal transcriptional cell types (clusters), grouped by GABAergic and glutamatergic neuronal classes, are shown.

**b**, Similar to (a), but quantification of E35a inclusion in Tasic 2018 dataset.

**c**, Quantification of E35a inclusion in different neuron types in adult mouse cortex and hippocampus.

**Extended Data Figure 3: Differential splicing of *Ank3* microexon E35a in diverse tissues.**

**a**, Quantification of E35a inclusion in different adult mouse tissues.

**b**, Quantification of E35a inclusion in different adult human tissues using GTEx RNA-seq data.

**Extended Data Figure 4: Dynamic *Ank3* E35a inclusion in different developmental stages and upon neuronal activation.**

**a,** Exon inclusion during the differentiation of mouse embryonic stem cells (mESCs) to glutamatergic excitatory neurons. DIV: days *in vitro*. Cells at DIV1-28 represent young and maturing neurons.

**b,** Exon inclusion during mouse cortex development.

**c,** Exon inclusion during human cortex development. pcw: post-conception weeks.

**d**, Changes of *Ank3* E35a inclusion (left) and Mbnl2 expression (right) in primary hippocampal neurons upon KCl treatment. FDR from differential splicing or expression analysis (derived by the Quantas pipeline) is indicated. The RNA-seq data used for splicing and gene expression quantification were obtained from Quesnel-Vallieres et al. 2016. This dataset was derived from cultured hippocampal cells dissociated from E16.5 mice, which were depolarized with 55 mM KCl for 3 h.

**e**, Similar to (d), but the RNA-seq data used for splicing and gene expression quantification were obtained from Ataman et al. 2016. This dataset was derived from cultured cortical cells dissociated from E16.5 mice, which were depolarized at DIV7 with 55 mM KCl for 6 h.

**Extended Data Figure 5: *Ank3* E35a encodes a peptide in an intrinsically disordered region between UPA domain and death domain.**

**a,** Alignments of experimentally determined AnkB domain structures (PDB: 4D8O) with AnkG domain structures predicted by AlphaFold3. Ankyrin repeats are not shown.

**b**, AlphaFold3-predicted AnkG domain structures with (left) or without (right) the peptide encoded E35a. The giant exon E37 is not included in the modeling. The death domain (DD) structures together with the upstream loop region with and without E35a are aligned (middle). Note the E35a-encoded peptide, indicated in red, is located in the intrinsically disordered region between the UPA and DD domains. Also, the inclusion of E35a converts the downstream 9aa-peptide from a loop into part of an extended alpha helix, highlighted in yellow.

**Extended Data Figure 6: Generation and validation of *Ank3* E35a^-/-^ mice.**

**a,** A schematic showing the generation of an *Ank3* E35a-deletion (*Ank3* E35a^-/-^) mouse model. A pair of single guide RNAs (sgRNAs) flanking the microexon E35a was used to target CRISPR-Cas9 to delete the microexon in the germline, which forces the generation of the exon-skipping isoform.

**b**, A schematic showing the design of single guide RNAs (sgRNAs, blue) for CRISPR-Cas9 gene editing. Forward/reverse primers for genotyping are indicated in red.

**c**, PCR genotyping of founder mice confirmed the deletion of the microexon (shaded samples).

**d**, RT-PCR validation of E35a splicing in adult wild-type (WT) and *Ank3* E35a^-/-^ cortex and cerebellum. Note that the exon is mostly skipped in the cortex but predominantly included in the cerebellum at the bulk tissue levels.

**e**, Body weight of WT and *Ank3* E35a^-/-^ littermate mice at 3 months old. Number of mice used for analysis: n = 15, 9 males and 6 females for WT; n = 14, 6 males and 8 females for mutant. No significance (p>0.05), Mann-Whitney *U* test.

**Extended Data Figure 7: Electrophysiological properties of *Ank3* E35a^-/-^ non-FS interneurons were largely unaffected.**

**a,** Two parameters used to categorize the *Dlx6a* interneurons into FS and non-FS interneurons: the membrane time constant (tau) and the number of APs elicited with a 400 pA current step at a holding potential of either -65 or -70 mV (the holding potential was chosen based on the nearest to the neuron RMP and with injected current lower than 100 pA). *Dlx6a* interneurons with a membrane time constant < 15 ms and > 35 APs (firing frequency of 70 Hz) were identified as FS (n = 42, shaded area). Neurons not following these criteria were considered non-FS (n = 52, white area), including putative somatostatin+ and vip+ interneurons.

**b**, Age distribution of patch clamp recordings for WT (8 mice, 48 cells) and *Ank3* E35a^-/-^ (8 mice, 46 cells) genotypes.

**c**, Average number of APs elicited by 500-ms current steps in FS interneurons, grouped by age into late-adolescent (P < 60) and adult (P > 60) mice. No age-related differences were detected in WT (left; two-way ANOVA, F_(1,19)_ = 0.1577, p = 0.6957) or Ank3 E35a⁻/⁻ mice (right; two-way ANOVA, F_(1,19)_ = 0.0547, p = 0.8176).

**d**, The fAHP was measured from the first 5 APs evoked by 500-ms current step at rheobase + 100 pA from RMP. No differences were observed between genotypes in either non-FS (left, two-way ANOVA, F_(1, 50)_ < 0.0005, p = 0.9809) or FS (right, two-way ANOVA, F_(1, 40)_ = 2.258, p = 0.1408) interneurons. **e**, Representative traces showing the initial firing response to a 500-ms current step at rheobase + 100pA from RMP in non-FS interneurons (left). From these traces, latency to the first AP (middle panel), and the first ISI (right panel) were quantified. Scale bar: 50 ms (horizontal).

**f,** Representative traces (left panel) from the firing pattern of WT and *Ank3* E35a^-/-^ non**-**FS interneuron at RMP in response to a 500-ms current stimulus with an amplitude of 200 pA. Scale bar: 40 mV (vertical). Input-output curves (right panel) show the relation between the number of APs and current step amplitude (two-way mixed ANOVA, F_(1, 50)_ = 0.9508, p = 0.3342). Average data are presented as mean ± SEM.

**g.** Rheobase from data in (f).

**h-j,** Passive membrane properties: RMP (h), input resistance (i) and capacitance (j) measured at RMP. **k**, The first AP evoked by the rheobase in panel (Fig. 3f in the main text) was used to measure the following properties in FS interneurons: voltage threshold, amplitude, width at half-height amplitude, maximum rise slope, and maximum decay slope. Scale bar: 40 mV.

**l**, The first AP evoked by the rheobase in panel (f) was used to measure the following properties in non-FS interneurons: voltage threshold, amplitude, width at half-height amplitude, maximum rise slope, and maximum decay slope.

**m**, Properties of the first AP at a common voltage (−70 mV) evoked from rheobase with 5 ms current injections in putative non-FS. The following properties of the first AP were analyzed: rheobase, voltage threshold, amplitude, width at half-width, maximum rise slope, and maximum decay slope. Data from 22 Non-FS interneurons (WT = 9, *Ank3* E35a^-/-^ = 13).

Mann Whitney U test was used in panels (b-d, g-j, k-m). Each dot represents data from a single cell. Data are presented as means ± SEM. Exact P-values are indicated in the graphs (p < 0.05 are in bold).

**Extended Data Figure 8: Analysis of AIS in cultured *Ank3* KO mouse hippocampal neurons transfected with AnkG 480kDa isoform with or without E35a.**

**a**, Schematics showing that hippocampal neurons from AG^E22/23-flox^ (*Ank3* KO) mice were cultured and transfected on DIV3 with plasmid expressing AnkG 480 kDa isoform with or without E35a.

**b**, On DIV7, neurons were fixed and immunostained against ankyrin-G and β4-spectrin (β4) and representative images are shown.

**c,d**, Intensity of AnkG (c) or β4-spectrin (d) in AIS are plotted against the distance from the soma. AIS lengths were quantified based on 8-10 cells per condition using an unpaired Student’s t-test.

Quantitative data are mean ± SEM. ns (p>0.05), not significant.

**Extended Data Figure 9: Behavioral analyses of WT and *Ank3* E35a^-/-^ mice.**

**a,** A schematic showing the timeline and sequence of different behavior assays.

**b**, Gait analysis of *Ank3* E35a^-/-^ and WT control mice. Left: A schematic of parameters analyzed is shown: fore paw-hind paw distance [FP-HP, Average (AB,CD)], step length [Average (AA,BB,CC,DD)], stance width [Average (AC, BD)] and alternation coefficient [(AB/BD)]. Right: Quantification in WT and *Ank3* E35a^-/-^ mice (mean ±SEM) is shown.

**c**, Beam walk test. A schematic of beam walk apparatus is shown at the top. The bar plots show quantification of Mean Velocity (cm/s), Mean Meander (deg/cm), Mean mobility, and Toe Slips in WT and *Ank3* E35a^-/-^ mice.

**d**, Elevated plus maze (EPM) test. A schematic of the EPM apparatus is shown at the top. The bar plots show the quantification of % Closed Arm Time, % Open Arm Time and Total Entries in WT and *Ank3* E35a^-/-^and mice.

**e**, Open field test. A schematic of open field apparatus is shown at the top. The bar plots show the Center Time (s), quantification of Distance (cm), and Vertical Activity (vertical counts) in WT and *Ank3* E35a^-/-^ mice.

**f**, Y-maze test. A schematic of Y-maze apparatus is shown at the top. The bar plots show the quantification of Spontaneous Activity and Total Entries in WT and *Ank3* E35a^-/-^ mice.

**g**, 3-chamber test of *Ank3* E35a^-/-^ and WT control mice. Left: A schematic of the 3-chamber test apparatus is shown. Right: Quantification in WT and *Ank3* E35a^-/-^ mice (mean ± SEM) is shown. Statistical analysis: For each panel, mean ± SEM is shown. Mann-Whitney *U* test is used except for open-field test, in which 2-way ANOVA with Repeated Measures was used. Intra-genotype comparisons in the 3-chamber test were performed using paired t-test. Number of 3 month-old mice used for 3-chamber test in panel (e): WT (n = 21; 11 males, 10 females); *Ank3* E35a-/- (n = 19; 9 males, 10 females). The number of 3 month-old mice used for other behavior tests in panels (b-d, f-h): WT (n = 15; 9 males, 6 females); *Ank3* E35a^-/-^ (n = 14; 6 males, 8 females). Only p < 0.05 are reported.

