## Supplementary fig for "A neuron type-specific microexon in *Ank3*/ankyrin-G modulates calcium activity and neuronal excitability"

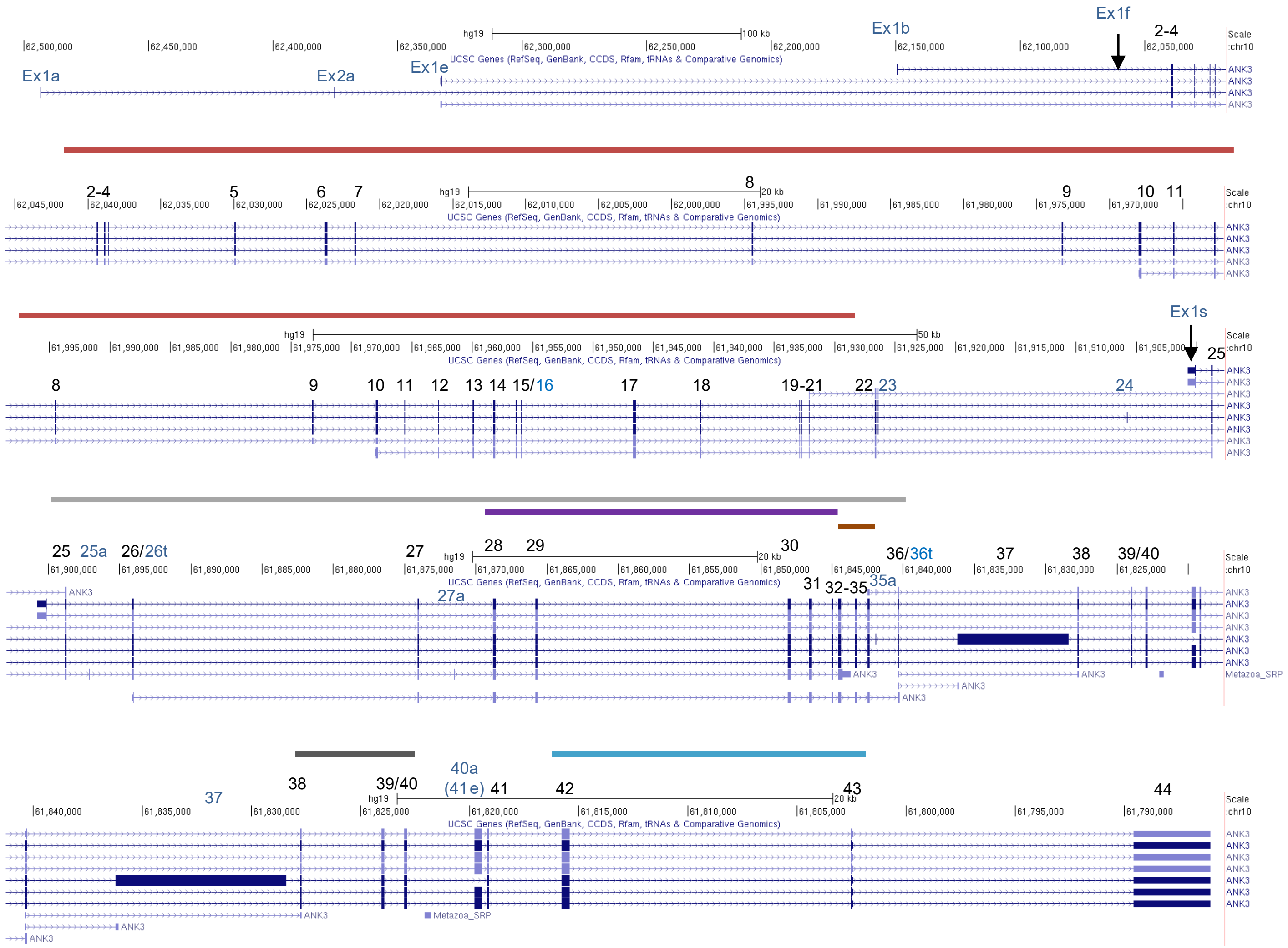


Exon 2-21: Membrane binding domain

Exon 25-36: Spectrin-binding domain
28-32: ZU5N/C
33-35: UPA

38-40: Death domain (DD)

40-43: C-terminal domain (CTD)

Extended Data Fig. 1

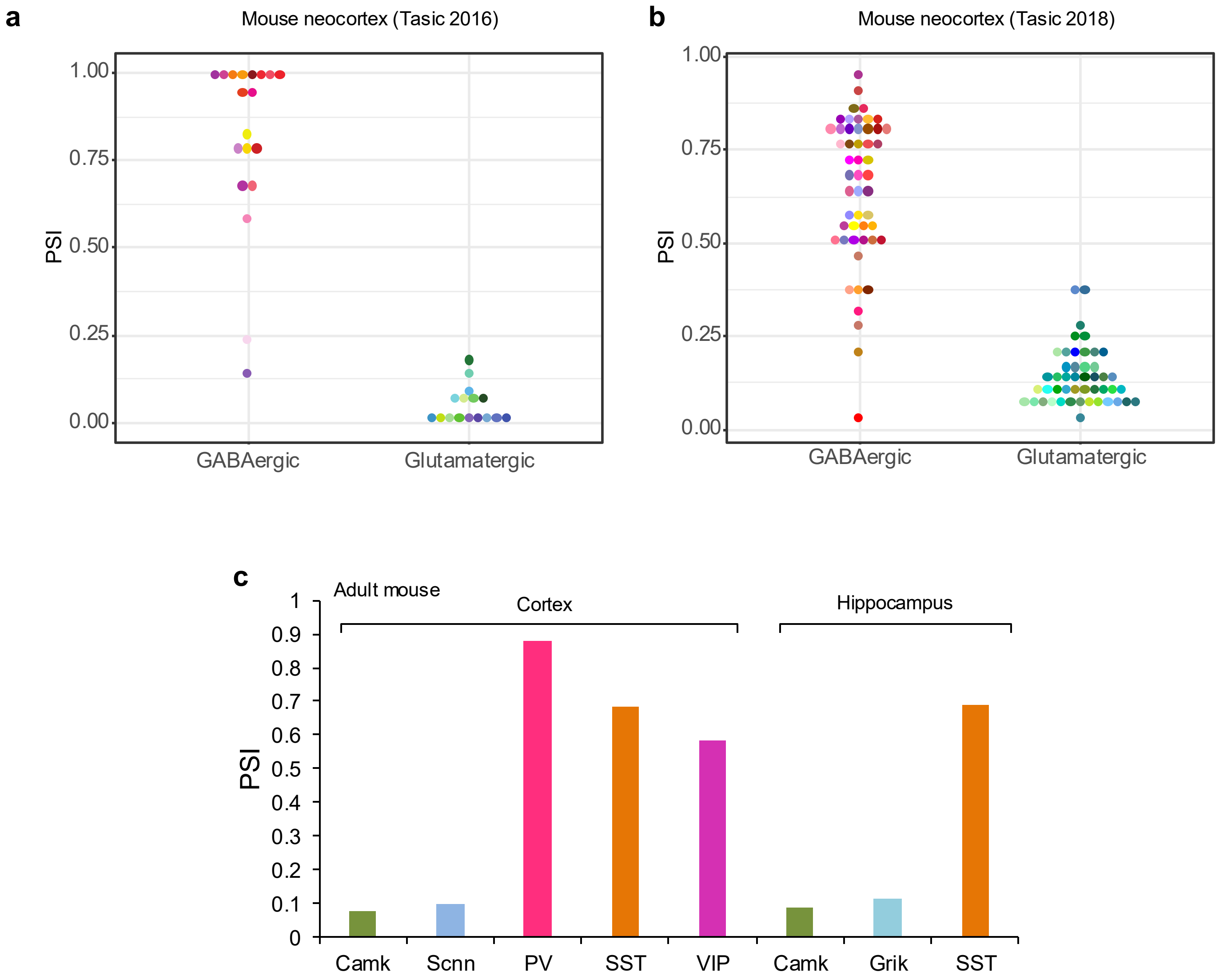


Extended Data Fig. 2

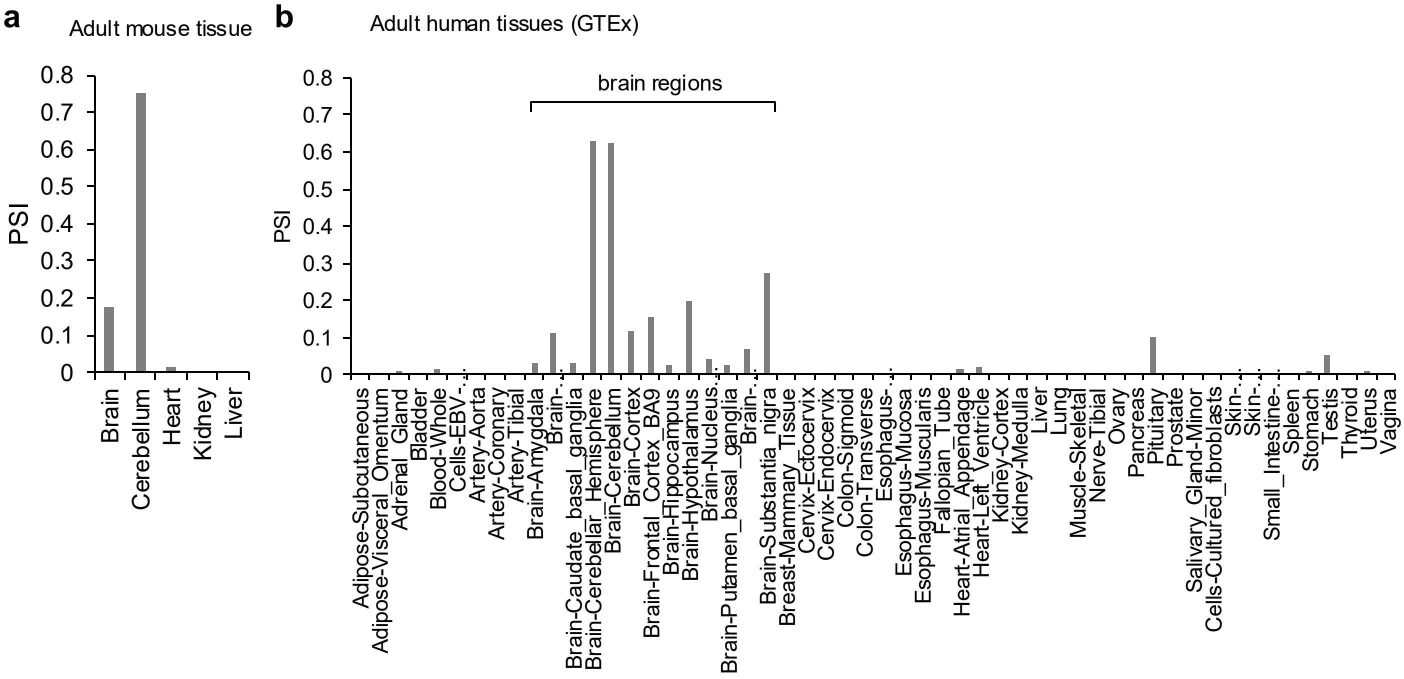


Extended Data Fig. 3

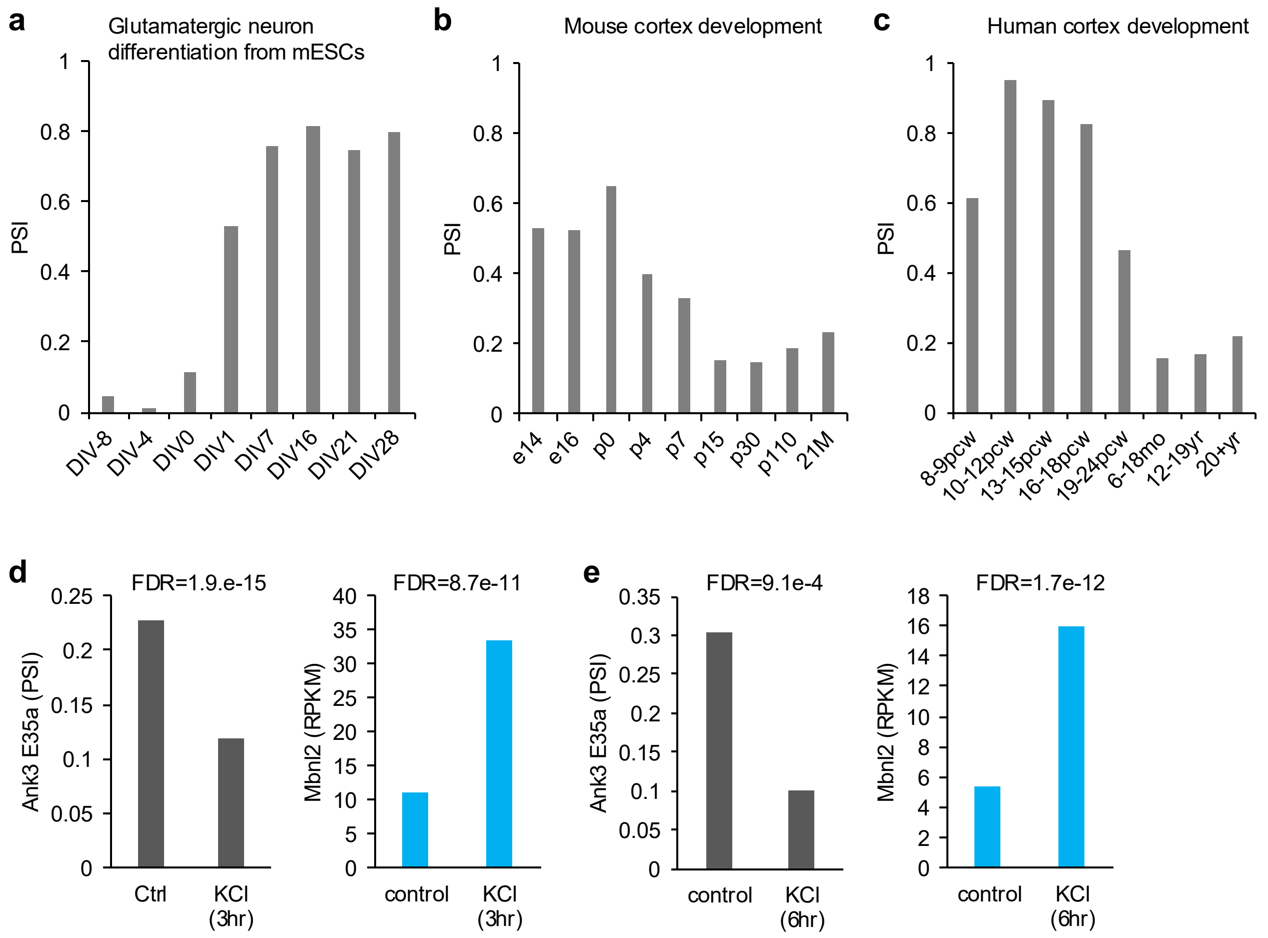


Extended Data Fig. 4

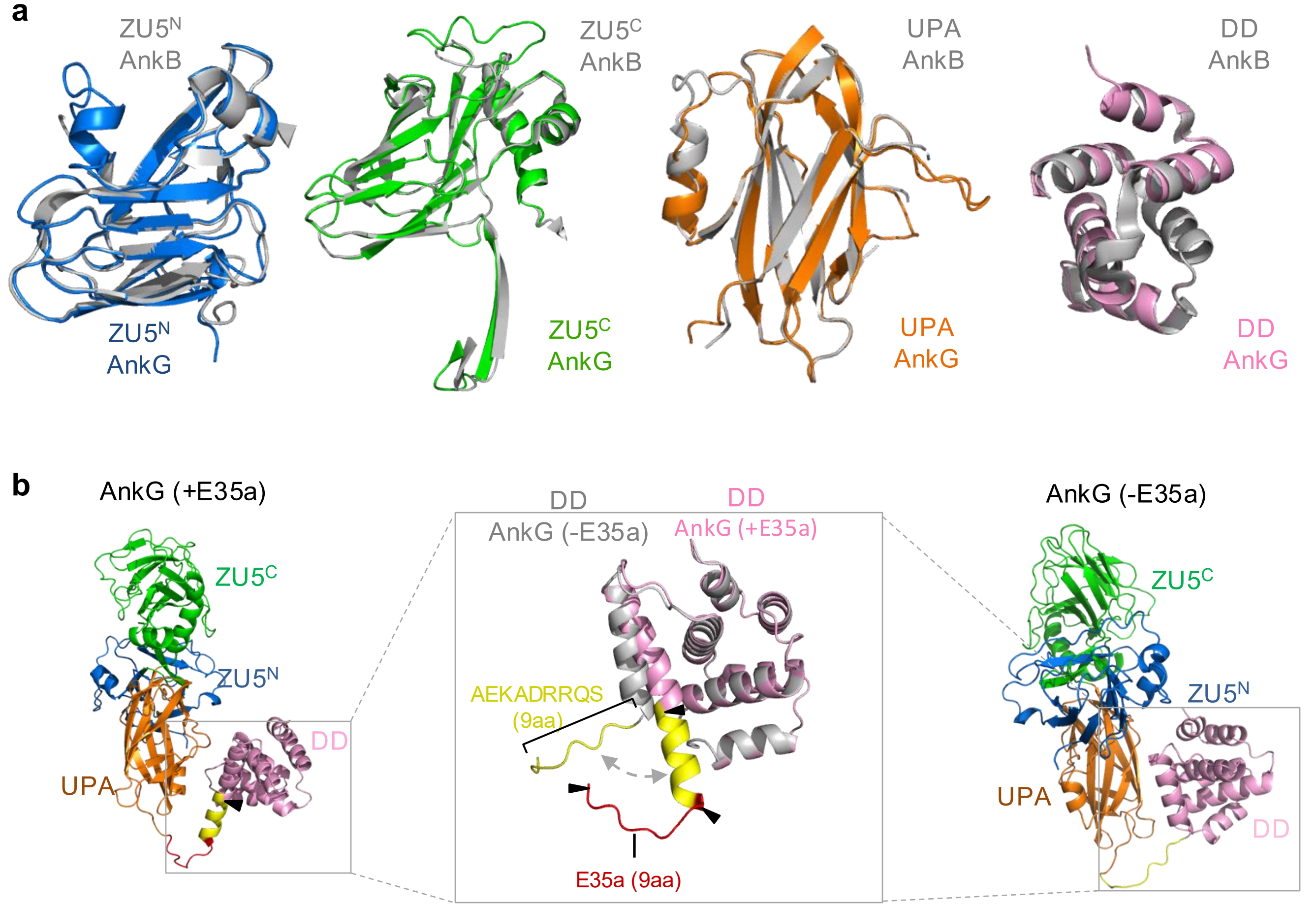


Extended Data Fig. 5

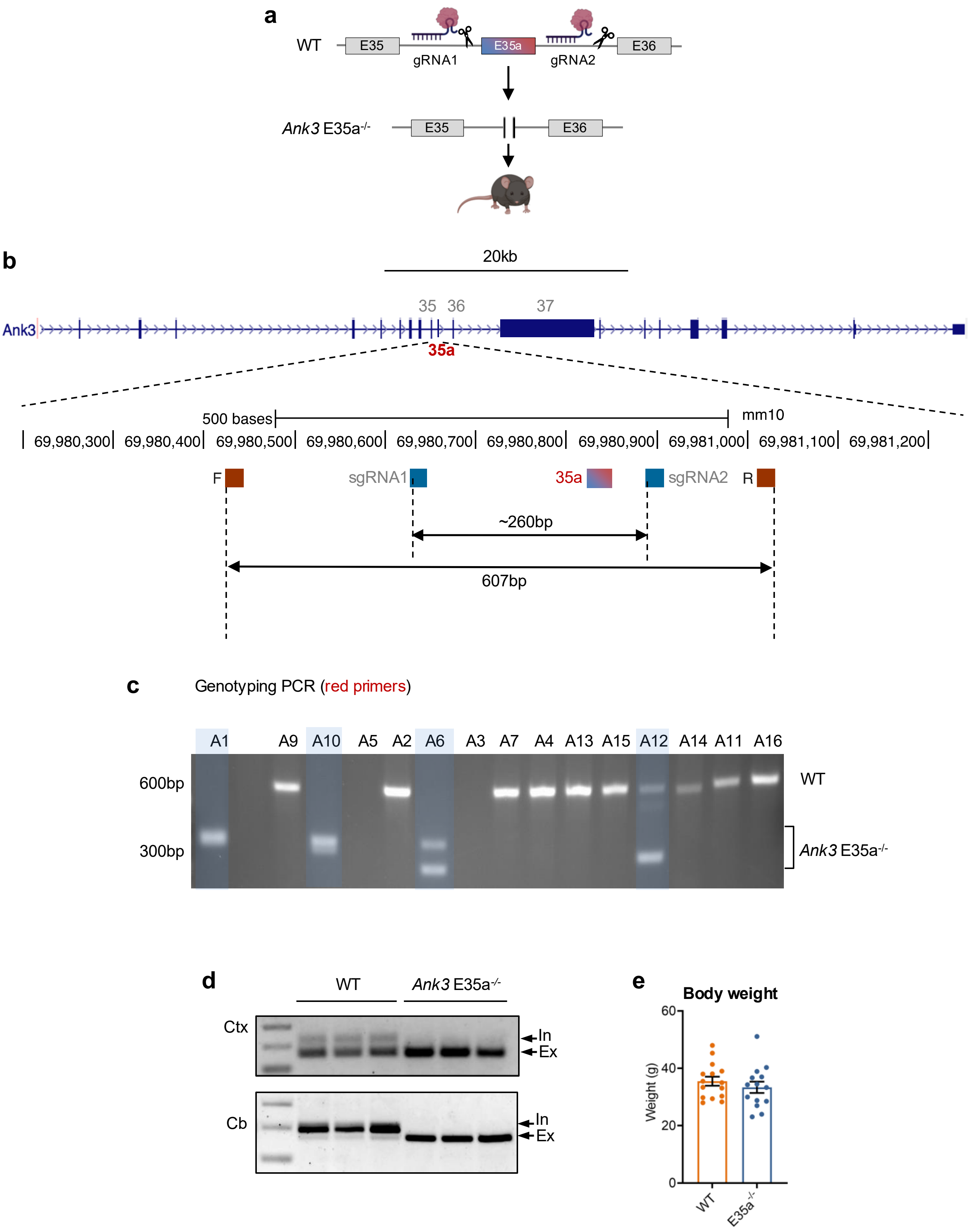


Extended Data Fig. 6

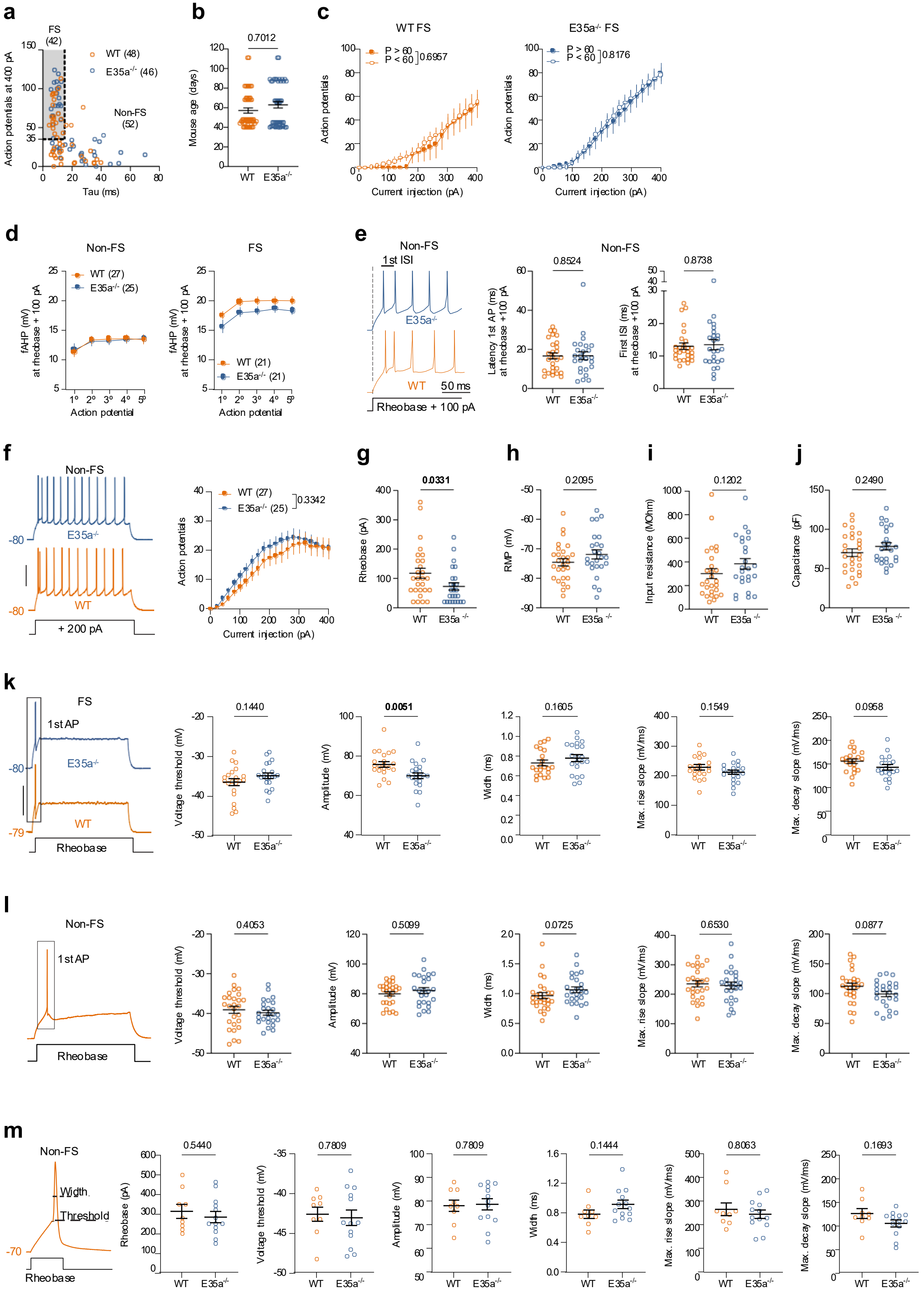


Extended Data Fig. 7

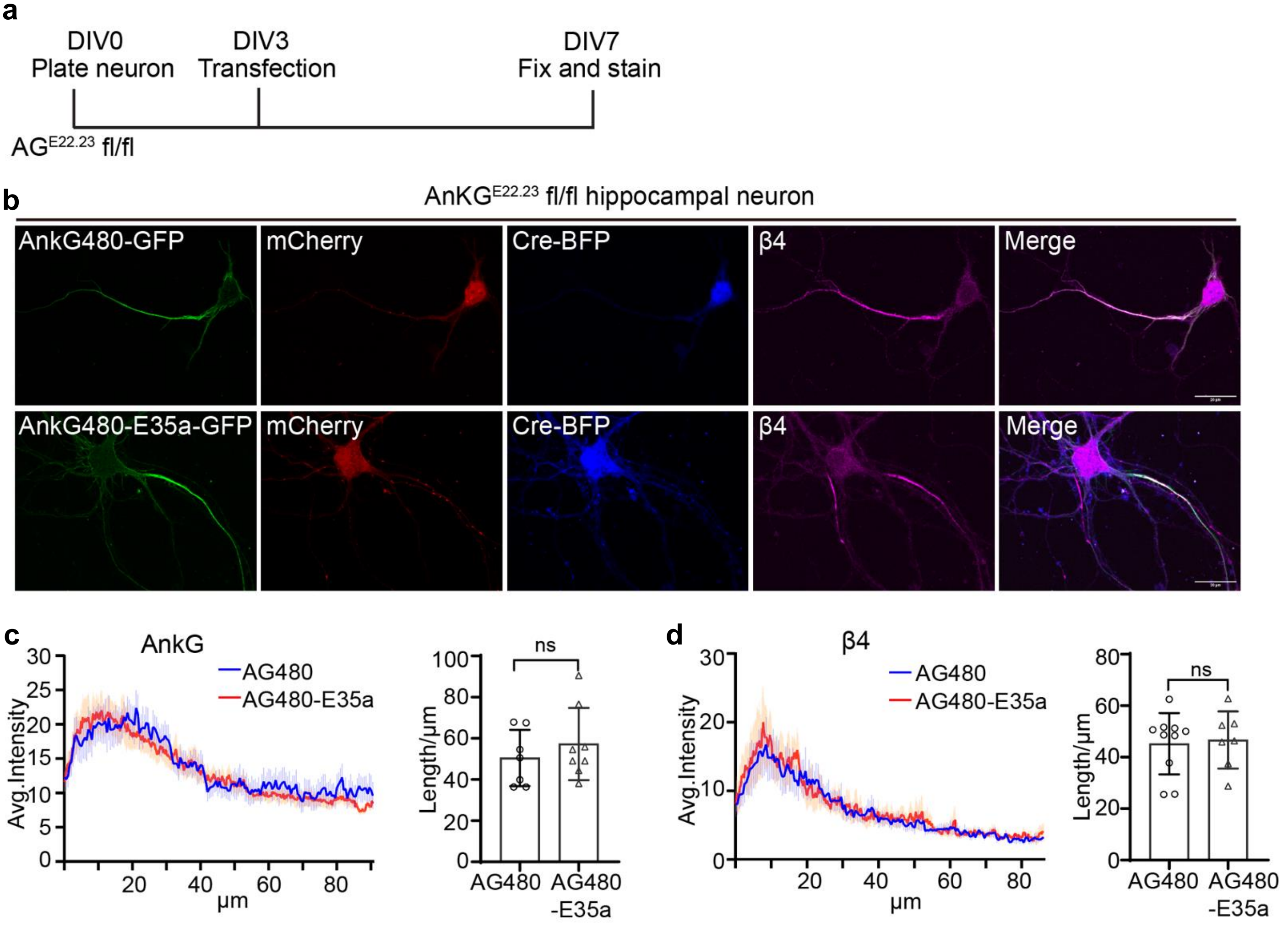


Extended Data Fig. 8

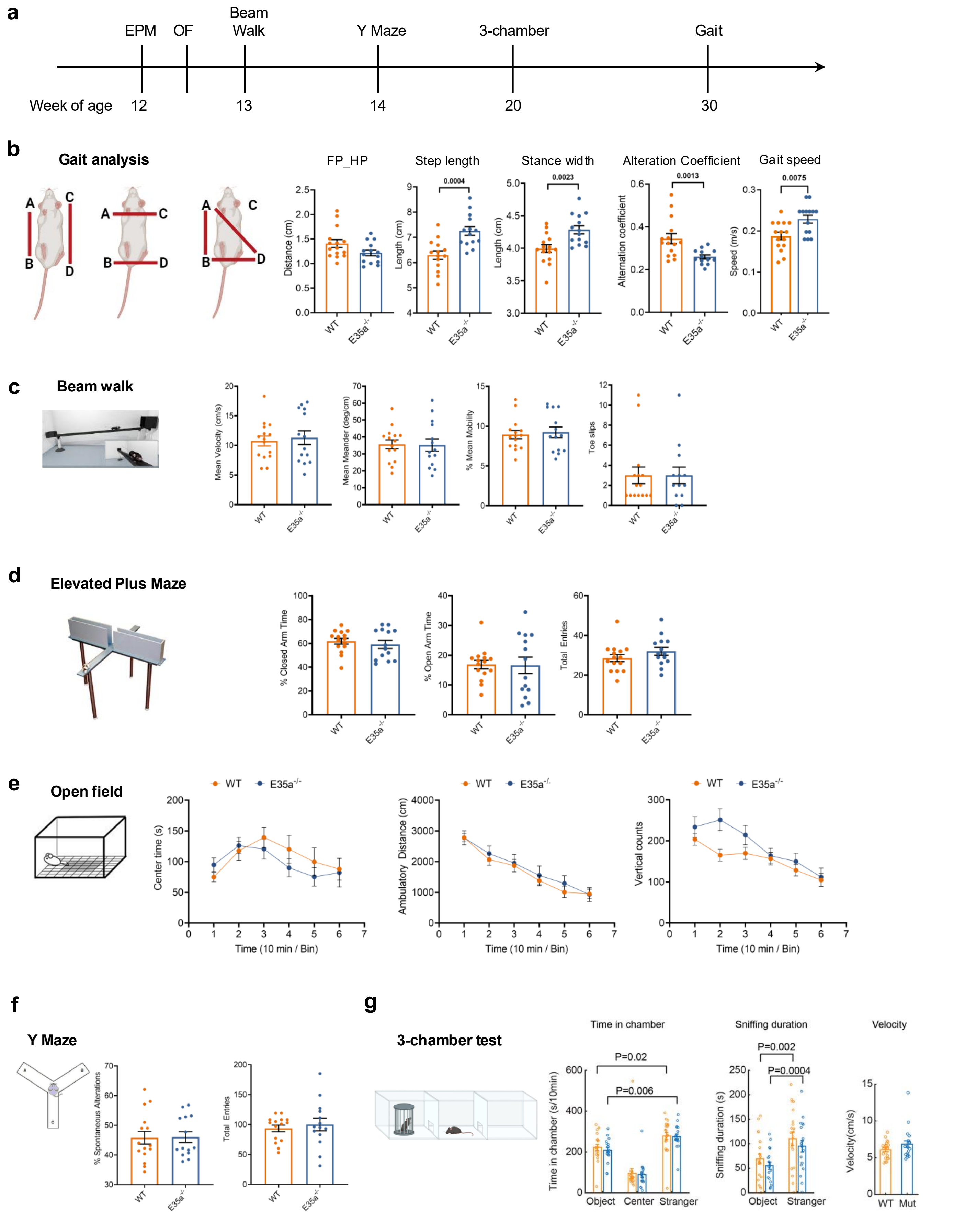


Extended Data Fig. 9
